# TACI Regulates Marginal Zone B Cell Development

**DOI:** 10.1101/2025.09.05.674478

**Authors:** Daisy H. Luff, Lesley Vanes, Stefan Boeing, Victor L. J. Tybulewicz

## Abstract

The mature B cell compartment consists of follicular (FO) and marginal zone (MZ) B cells, which develop from transitional type 2 (T2) B cells and mount T-dependent and T-independent antibody responses, respectively. TACI, a member of the TNF receptor superfamily, is expressed on all mature B cells, with highest levels on MZ B cells and plasma cells. Previous studies reported that TACI is a negative regulator of B cell survival. However, this conclusion is confounded by elevated levels of BAFF, a cytokine that supports B cell survival, in TACI- deficient mice. We now show that TACI does not directly regulate B cell survival but rather has a cell-intrinsic role in MZ B cell development. Loss of TACI leads to reduced MZ B cell numbers and impaired T-independent antibody responses. Mechanistically, we show that TACI is required for MZ B cell development from T2 B cell precursors via activation of the PI3K-AKT pathway and subsequent inhibition of the FOXO1 transcription factor.

## Introduction

B lymphocyte development is a continuous process in adult mammals, starting in the bone marrow where B cell progenitors undergo immunoglobulin heavy chain and then light chain gene rearrangements (Hardy and Hayakawa, 2001). Successful rearrangement results in the production of immature B cells expressing a diverse repertoire of B cell antigen receptors (BCRs) in the form of surface immunoglobulin M (IgM) associated with CD79A and CD79B signaling subunits. These immature B cells exit the bone marrow and migrate through the blood to the spleen, arriving as transitional type 1 (T1) B cells that mature to T2 B cells and finally into either follicular (FO) or marginal zone (MZ) mature B cells, which can mount T-dependent and T-independent antibody responses, respectively (Pillai and Cariappa, 2009). FO B cells continuously recirculate between the spleen and other lymphoid organs and reside in the follicles of these organs. In contrast, MZ B cells are enriched in the marginal zone of the spleen where they rapidly respond to blood-borne antigens and capture and deliver them to the follicle (Cinamon et al., 2008). A third type of mature B cell, B1 cells, are generated predominantly during gestation in the fetal liver, have a more limited BCR repertoire, are maintained by self-renewal and localize to both lymphoid as well as non-lymphoid tissues such as peritoneal and pleural cavities (Hardy, 2006).

The development of T2 B cells into mature B cells and the decision whether they become FO or MZ B cells is incompletely understood. It is regulated in part by BCR signaling strength, which is dependent on reactivity of the BCR to self-antigens (Tan et al., 2019). Too little BCR signaling and cells die by neglect, whereas too much auto-reactivity and BCR signaling leads to elimination of the B cells by negative selection. In between these two outcomes, T2 B cells are positively selected into mature B cells, with the FO fate being acquired by cells with a broad range of self-reactivity and thus BCR signaling strength, whereas the MZ fate is adopted by B cells with stronger BCR signaling resulting from a higher and narrower range of self-reactivity (Noviski et al., 2019). In addition to the BCR, MZ B cell development also requires signaling from the NOTCH2 receptor (Pillai and Cariappa, 2009).

The overall size of the FO and MZ B cell pools remains stable in the adult animal (Cancro, 2004). Both FO and MZ B cells are unable to self-renew and are maintained by continuous production of immature B cells in the bone marrow. Homeostasis is determined by a balance between the rate of entry of B cells differentiating from T2 B cells into the mature FO and MZ B cell pools and the rate of exit of cells from these pools because of death or activation-induced differentiation into germinal center B cells, memory B cells or plasma cells. Thus, the regulation of B cell survival is critical for the maintenance of mature B cell pool sizes.

Two receptors are known to regulate the survival of mature B cells - the BCR and BAFFR (Schweighoffer and Tybulewicz, 2018). Several studies suggest that the BCR delivers a survival signal via the associated CD79A and CD79B proteins and the SYK protein tyrosine kinase (Kraus et al., 2004; Lam et al., 1997; Schweighoffer et al., 2013). Inducible genetic ablation of IgM, CD79A or SYK results in the loss of all mature B cells (FO, MZ and B1 cells). This ’tonic’ BCR survival signal is below the level required for B cell activation and may be dependent on binding to self-antigens. BAFFR, a member of the TNF receptor superfamily, is a receptor for the cytokine BAFF. Genetic ablation of either BAFF or BAFFR or antibody-mediated blockade of BAFF-BAFFR binding results in loss of T2 B cells as well as FO and MZ mature B cells, but not B1 cells (Gross et al., 2001; Gross et al., 2000; Mackay et al., 2010; Mackay et al., 1999; Miller and Hayes, 1991; Rauch et al., 2009; Sasaki et al., 2004; Schiemann et al., 2001; Schneider et al., 2001; Shulga-Morskaya et al., 2004; Smulski and Eibel, 2018; Thompson et al., 2001). Despite the importance of BAFF-BAFFR signaling for B cell survival, the biochemical mechanism by which the receptor delivers survival signals is unclear. The best described signaling pathway from BAFFR involves BAFF-dependent binding of TRAF3 to BAFFR resulting in the activation of IKK1 and the non-canonical NF-κB pathway (Gardam and Brink, 2014). However, inducible deletion of *Ikk1* had no impact on the numbers of mature B cells, demonstrating that this pathway was not required for mature B cell survival (Jellusova et al., 2013). Interestingly, BAFFR has been shown to transduce survival signals in part via BCR, CD19 and SYK, leading to activation of phosphoinositide 3-kinase (PI3K) and ERK pathways (Jellusova et al., 2013; Keppler et al., 2015; Schweighoffer et al., 2013). However, it is not known how BAFFR signals are transduced to these proteins, since TRAF3, the only protein reported to bind to the intracellular domain of BAFFR, has not been shown to transduce signals to the BCR, SYK, ERK or PI3K pathways.

To investigate the mechanism by which BAFFR signals, we used affinity purification of BAFFR and mass spectrometry to identify new proteins that interact with BAFFR. We show that BAFFR associates with TACI, a related TNF-receptor superfamily protein that binds BAFF as well as the cytokine APRIL. Previous studies had concluded that TACI was a negative regulator of B cell survival, since TACI-deficient mice have expanded numbers of B cells (Shulga-Morskaya et al., 2004; von Bulow et al., 2001; Yan et al., 2001). However, we now demonstrate that TACI does not directly regulate the survival of either FO or MZ B cells. Instead, we show for the first time that TACI deficiency leads to a loss of MZ B cells, owing to a cell-intrinsic function in the development of MZ B cells from T2 B cells. Furthermore, we show that TACI regulates MZ development most likely by transducing signals via PI3K and AKT, leading to inhibition of the FOXO1 transcription factor and reduced *Klf2* expression. Thus, our study uncovers an unexpected role for TACI and reveals a novel pathway regulating the MZ versus FO cell fate decision, functioning alongside the BCR and NOTCH2. The work also widens our view of how TNF-receptor superfamily members regulate the development and maintenance of the mature B cell repertoire.

## Results

### Loss of TACI leads to reduced numbers of marginal zone B cells

To gain a better understanding of how BAFFR transduces signals leading to B cell survival, we generated mice expressing BAFFR with two copies of a StrepTag II affinity tag at its C-terminus (Fig. S1A). We used affinity purification to isolate BAFFR-TwinStrepTag from LPS-activated B cells, which had been untreated or stimulated with BAFF for 15 min, followed by mass spectrometry to identify proteins associated with BAFFR. As expected, we identified BAFFR in all pull-downs and BAFF in pull-downs from stimulated cells. One other protein, the receptor TACI, co-purified with BAFFR and this association only occurred in BAFF-stimulated cells (Fig. S1B, Table S1). Thus, we hypothesized that TACI may co-operate with BAFFR to regulate B cell survival. In considering this idea, we noted previous publications showing that loss of TACI (encoded by the *Tnfrsf13b* gene) in *Tnfrsf13b*^-/-^ mice results in increased B cell numbers, suggesting that TACI is a negative regulator of B cell survival (Shulga-Morskaya et al., 2004; von Bulow et al., 2001; Yan et al., 2001). However, this conclusion is confounded by the observation that *Tnfrsf13b*^-/-^ mice have elevated levels of BAFF in the serum (Bossen et al., 2008; Sintes et al., 2017). Since overexpression of BAFF leads to increased B cells (Mackay et al., 1999), the higher B cell numbers in TACI-deficient mice could be caused by elevated BAFF in the serum.

To extend these studies, we analyzed TACI-deficient mice and confirmed that they have a 3- fold increase in circulating levels of BAFF and increased numbers of both FO and MZ B cells in the spleen, although the increase is larger for FO B cells resulting in a decrease in the fraction of mature B cells that are MZ B cells (Fig. 1A, S1C, D). In agreement with increased BAFF-induced signaling through BAFFR, TACI-deficient FO B cells had elevated levels of ICOSL, a known target of BAFFR signaling (Fig. S1E) (Ou et al., 2012).

**Figure 1.**
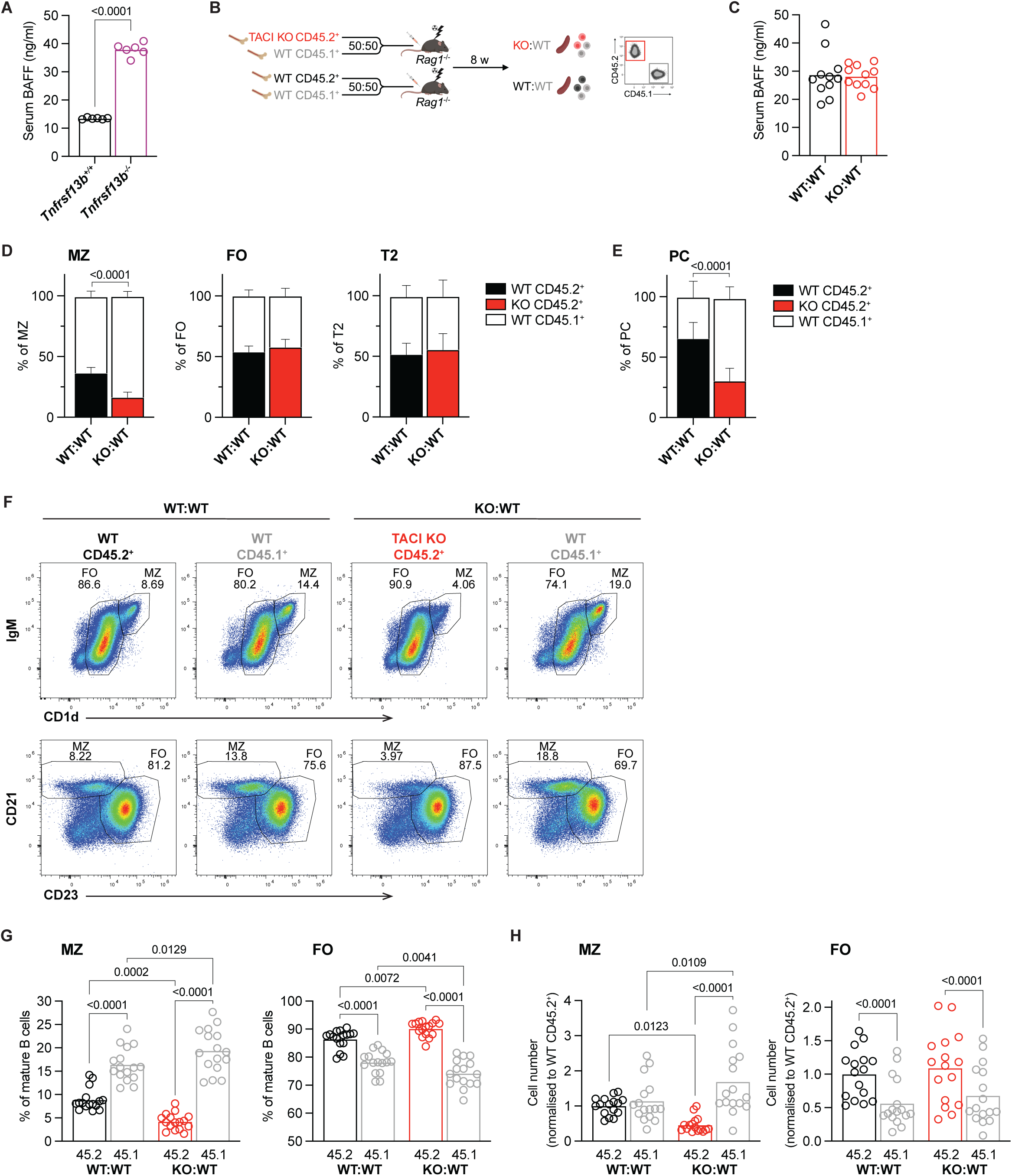
TACI deficiency leads to reduced MZ B cells. **A.** Concentration of BAFF in the serum of *Tnfrsf13b*^+/+^ and *Tnfrsf13b*^-/-^ mice. Bars show the mean. Each point represents one mouse. **B.** Mixed bone marrow chimera generation. Bone marrow from *Tnfrsf13b*^+/+^ or *Tnfrsf13b*^-/-^ littermates (expressing the CD45.2^+^ allele) was mixed 50:50 with bone marrow from wildtype CD45.1^+^ mice then injected into sub-lethally irradiated *Rag1*^-/-^ mice. B cells were analyzed at least 8 weeks later. **C.** Concentration of BAFF in the serum of WT:WT and KO:WT chimeras. Bars show the mean. Each point represents one mouse. **D.** Stacked bar charts showing the percentages of CD45.2^+^ and CD45.1^+^ cells within the splenic MZ (CD93^-^CD1d^hi^IgM^hi^), FO (CD93^-^CD1d^med^IgM^+^) and T2 (CD93^+^IgD^+^IgM^+^) B cell (B220^+^CD19^+^) populations in WT:WT and KO:WT chimeras. Cells were analyzed by flow cytometry using the gating strategy in Fig. S2A. Bars show the mean ±95% CI. **E.** Stacked bar chart showing the percentages of CD45.2^+^ and CD45.1^+^ cells within the splenic plasma cell population (PC; TCRβ^-^CD138^+^CD19^lo^) in WT:WT and KO:WT chimeras. Bars show the mean ±95% CI. **F.** Representative flow cytometry plots showing the proportions of MZ and FO B cells within the CD45.2^+^ and CD45.1^+^ mature splenic B cell populations (CD93^-^B220^+^CD19^+^CD138^-^) in WT:WT and KO:WT chimeras. MZ and FO cells are gated via two different approaches. Numbers indicate the percentage of cells in that gate. **G, H.** Proportions (G) and numbers (H) of MZ (CD1d^hi^IgM^hi^) and FO (CD1d^med^IgM^+^) cells within the CD45.2^+^ and CD45.1^+^ mature splenic B cell populations (CD93^-^B220^+^CD19^+^CD138^-^) in WT:WT and KO:WT chimeras. Bars show the mean. Each point represents one mouse. Black: WT CD45.2^+^; red: TACI KO CD45.2^+^; grey: WT CD45.1^+^. Statistical tests: Welch’s t test (A, C), two-way ANOVA with Fisher’s LSD test (D, E), repeated measures two-way ANOVA with Fisher’s LSD test (G, H). Numbers above graphs indicate p- values where p≤0.05 otherwise no value is shown. Sample numbers: 6 (A, D), 11 (C), 5 (E), 16 (G, H). Data pooled from 2 experiments representative of 4 experiments (D), 1 out of 2 experiments (E), pooled from 4 independent experiments (G, H) or from 6 independent experiments (A, D).

In view of the increased levels of BAFF, BAFFR signaling and mature B cell numbers in TACI- deficient mice, it is not possible to analyze the cell-intrinsic role of TACI by comparing B cells from TACI-deficient and wildtype (WT) mice. To resolve this issue, we generated mixed bone marrow chimeras, reconstituting irradiated RAG1-deficient mice with bone marrow from either *Tnfrsf13b*^-/-^ or *Tnfrsf13b*^+/+^ mice expressing CD45.2 (CD45.2^+^), mixed 50:50 with WT bone marrow from CD45.1^+^ mice, to generate KO:WT and WT:WT chimeras, respectively (Fig. 1B). Both sets of chimeras had similar levels of serum BAFF, confirming that this approach eliminates the confounding factor of elevated BAFF (Fig. 1C). In line with this, KO:WT chimeras did not display splenic B cell hyperplasia and TACI-deficient FO B cells from KO:WT chimeras had no change in ICOSL surface expression, consistent with unaltered BAFFR signaling (Fig. S2A-C). These observations indicate that the B cell hyperplasia and increased ICOSL expression seen in *Tnfrsf13b*^-/-^ mice are not cell-intrinsic effects of TACI loss, but more likely a consequence of increased levels of BAFF in the serum, and demonstrate that the mixed chimeras can be used to study the cell-intrinsic consequence of TACI deficiency.

Comparison of WT and TACI-deficient B cells taken from WT:WT and KO:WT chimeric mice showed that low levels of TACI expression can be detected on T1 and T2 B cells, with upregulation of TACI occurring as the cells mature into FO and MZ B cells (Fig. S2D). MZ B cells express substantially more TACI compared to FO B cells, and even higher levels are seen on plasma cells.

Surprisingly, further analysis of the chimeras showed that the proportion of TACI-deficient CD45.2^+^ MZ B cells was reduced in KO:WT chimeras compared to WT CD45.2^+^ MZ B cells in WT:WT chimeras (Fig. 1D, S2A). In contrast, loss of TACI had no effect on the proportions of FO B cells or T2 B cells, the direct precursors of MZ and FO B cells. Analysis of splenic plasma cells showed that the proportion and number of TACI-deficient plasma cells was reduced (Fig. 1E, S2A, S2E), in line with previous studies reporting the importance of TACI in plasma cell survival and differentiation (Eslami et al., 2024; Ou et al., 2012; Sintes et al., 2017; Tsuji et al., 2011). Furthermore, within the TACI-deficient mature B cells, the proportion and absolute number of MZ B cells was reduced, while the proportion of FO B cells was increased, but the absolute number of FO B cells was not significantly changed (Fig. 1F-H). Finally, loss of TACI had no effect on the numbers of T2 B cells (Fig. S2F). These observations demonstrate that loss of TACI results in a cell-intrinsic reduction in the number of MZ B cells, with no effect on the numbers of FO or T2 B cells.

### TACI is required for T-independent antibody responses

Given that MZ B cells are a major contributor to T-independent antibody responses (Li et al., 2022; Martin et al., 2001; Swanson et al., 2010), we examined whether the reduction in TACI- deficient MZ B cells in KO:WT chimeras would impair this immune response. To do this, we reconstituted RAG1-deficient mice with bone marrow from either *Tnfrsf13b*^-/-^ or *Tnfrsf13b*^+/+^ C57BL/6 (B6) mice, this time mixed 50:50 with WT bone marrow from 129S7 (129) mice, generating KO:wt and WT:wt chimeras, respectively (Fig. 2A). In these mixed chimeras, antibodies and B cells of B6 and 129 origins can be distinguished by IgH^b^ and IgH^a^ allotypes and by expression of Ly9.2 and Ly9.1, respectively. Once again, we found that the proportion and number of TACI-deficient MZ B cells was reduced in KO:wt chimeras compared to WT MZ B cells in WT:wt chimeras, whereas FO B cells were not affected (Fig. 2B-E, S2G). We immunized the chimeras with TNP-Ficoll, an antigen that stimulates a T-independent antibody response, and 7 days later measured serum levels of TNP-specific IgM and IgG1 of both a and b allotypes (Fig. 2A). We found that KO:wt chimeras had a greatly impaired IgM^b^ response compared to WT:wt chimeras, and a reduced IgG1^b^ response, whereas IgM^a^ and IgG1^a^ responses in the same mice originating from the 129-derived WT B cells were unaffected (Fig. 2F). In line with this, we observed a reduced expansion of TACI-deficient plasma cells compared to WT plasma cells in response to immunization with TNP-Ficoll, and no expansion of TACI-deficient plasmablasts (Fig. 2G-H, S2G). Together, these results suggest that the reduction in MZ B cells due to the absence of TACI leads to an impaired T-independent antibody response.

**Figure 2.**
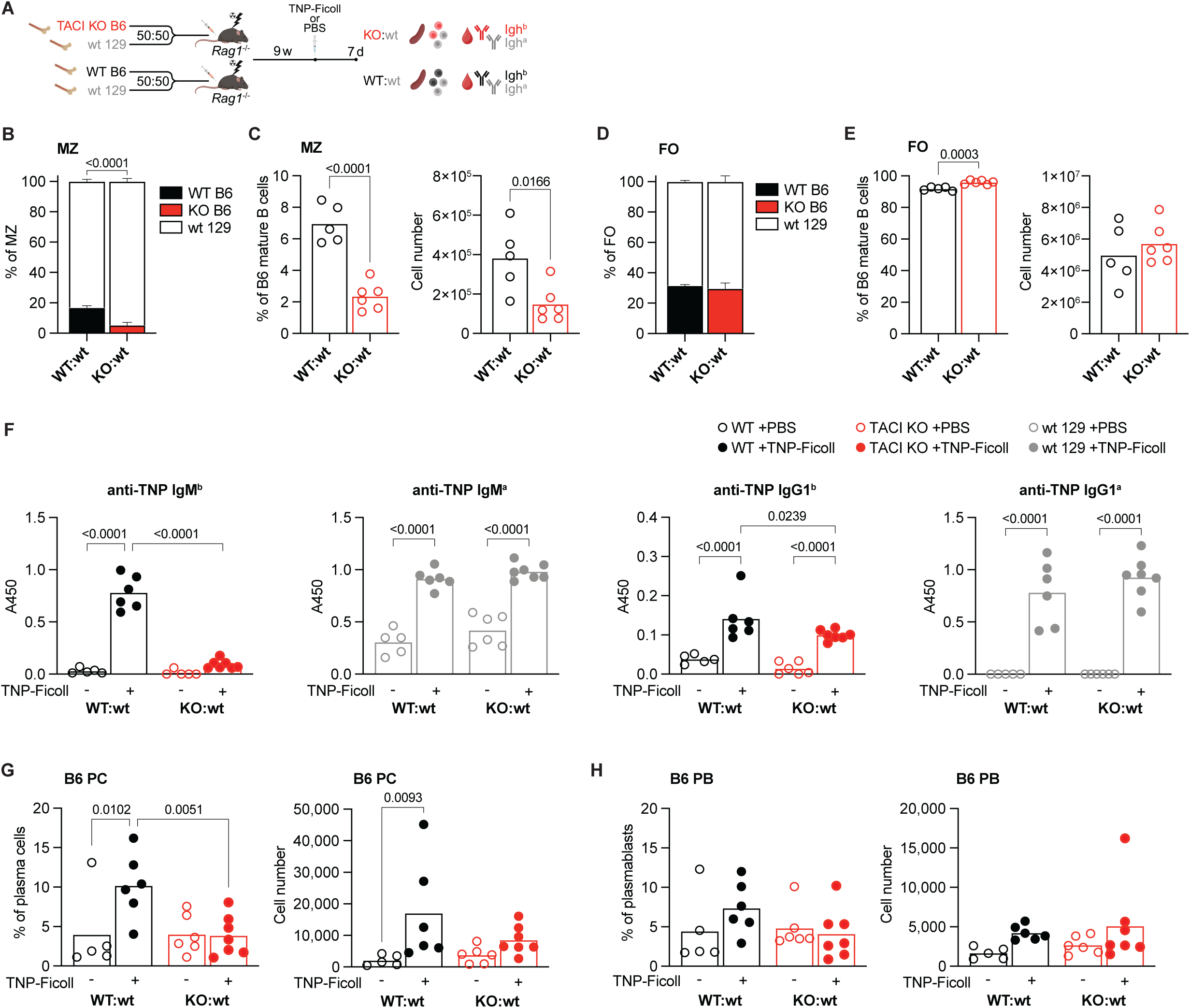
TACI is required for T-independent antibody responses. **A.** Mixed bone marrow chimera generation for studies of T-independent antibody responses. Bone marrow from *Tnfrsf13b*^+/+^ or *Tnfrsf13b*^-/-^ littermates on the C57BL/6 (B6) background (Ly9.1^-^) was mixed 50:50 with bone marrow from wildtype 129S7 (129) mice (Ly9.1^+^) then injected into sub-lethally irradiated *Rag1*^-/-^ mice, to generate WT:wt and KO:wt chimeras, respectively. 9 weeks later mice were injected i.p. with TNP-Ficoll or PBS as control. 7 days post-immunization, spleens and blood were harvested for analysis. Antibodies produced by C57BL/6 or 129S7 cells are of the Igh^b^ or Igh^a^ allotype, respectively. **B.** Stacked bar chart showing the percentages of B6 and 129 cells within the total splenic MZ B cell populations (B220^+^CD19^+^CD93^-^CD1d^hi^IgM^hi^) in WT:wt and KO:wt chimeras. Cells were analyzed by flow cytometry using the gating strategy shown in Fig. S2G. Bars show the mean ±95% CI. **C.** Proportion of MZ B cells (CD1d^hi^IgM^b^ ^hi^) within the B6 mature splenic B cell populations (Ly9.1^-^CD93^-^B220^+^CD19^+^CD138^-^) and cell numbers in WT:wt and KO:wt chimeras. Bars show the mean. Each point represents one mouse. Black: WT B6; red: TACI KO B6. **D.** Stacked bar chart showing the percentages of B6 and 129 cells within the total splenic FO B cell populations (B220^+^CD19^+^CD93^-^CD1d^med^IgM^+^) in WT:wt and KO:wt chimeras. Cells were analyzed by flow cytometry using the gating strategy shown in Fig. S2G. Bars show the mean +95% CI. **E.** Proportion of FO B cells (CD1d^med^IgM^b+^) within the B6 mature splenic B cell populations (Ly9.1^-^CD93^-^B220^+^CD19^+^CD138^-^) and cell numbers in WT:wt and KO:wt chimeras. Bars show the mean. Each point represents one mouse. Black: WT B6; red: TACI KO B6. **F.** TNP-specific IgM^b^, IgM^a^, IgG1^b^ and IgG1^a^ antibody responses in WT:wt and KO:wt chimeras 7 days after immunization with TNP-Ficoll or PBS control. IgM^b^ and IgG1^b^ originates from B6 B cells (black: WT; red: TACI KO), whereas IgM^a^ and IgG1^a^ originates from WT 129 B cells (grey). Antibody levels in serum were determined by ELISA and measured as absorbance at 450 nm (A450). Bars show the mean. Each point represents one mouse. **G, H.** Percentage and numbers of B6 cells (Ly9.1^-^) within the splenic plasma cells (PC; TCRβ^-^ CD138^+^B220^lo^CD19^int-lo^) (G) or plasmablasts (PB; TCRβ^-^CD138^+^B220^hi^CD19^int^) (H) in WT:wt and KO:wt chimeras 7 days after immunization with TNP-Ficoll or PBS control. Bars show the mean. Each point represents one mouse. Black: WT B6; red: TACI KO B6. Statistical tests: unpaired t test (C, E), two-way ANOVA with Fisher’s LSD test (B, D, F-H). Numbers above graphs indicate p-values where p≤0.05 otherwise no value is shown. Sample numbers: 5 (B-E, WT:wt; F-H, WT:wt +PBS; F, KO:wt +PBS IgM^b^ only), 6 (B-E, KO:wt; F-H, WT:wt +TNP-Ficoll, KO:wt +PBS except for IgM^b^ in F), 7 (F-H, KO:wt +TNP-Ficoll).

### TACI-deficient MZ B cells do not have a survival or homing defect

The reduction in splenic MZ B cells in the absence of TACI could be due to impaired development, homing or survival. In view of the association of TACI with BAFFR we hypothesized that TACI may regulate MZ B cell survival. To address this, we measured the turnover of MZ B cells in the mixed chimeras by administering EdU to the mice continuously for 6 weeks (Fig. 3A). In the steady state, only newly generated pro- and pre-B cells in the bone marrow proliferate and incorporate EdU into their DNA. In agreement with this, treatment of the mice with EdU for just 4 h showed no labelling of immature or mature splenic B cell populations of either genotype, confirming that they were not dividing (Fig. 3B, S3A-B). Thus, the accumulation of EdU^+^ immature and mature B cells measures influx of newly generated EdU^+^ B cells into the peripheral B cell pools and, given that these pools are maintained at a constant size, the rate of EdU^+^ cell entry is also a measure of the rate of efflux by death or differentiation. We found that the accumulation of EdU^+^ B cells in the immature T1, T2 and T3 B cell pools and the mature MZ and FO B cell pools was not affected by loss of TACI, demonstrating that the efflux of TACI-deficient MZ B cells from the MZ pool was unaltered and implying that TACI-deficient MZ B cells do not have a survival defect (Fig. 3B, S3A-B). To extend this, we transferred splenic B cells from mixed chimeras into CD45.1^+^CD45.2^+^ recipient mice and analyzed the mice 7 days later (Fig. 3C, S3C). We found no differences in the recovery of TACI-deficient MZ or FO B cells compared to WT cells, supporting our conclusion that loss of TACI does not affect MZ B cell survival (Fig. 3D).

**Figure 3.**
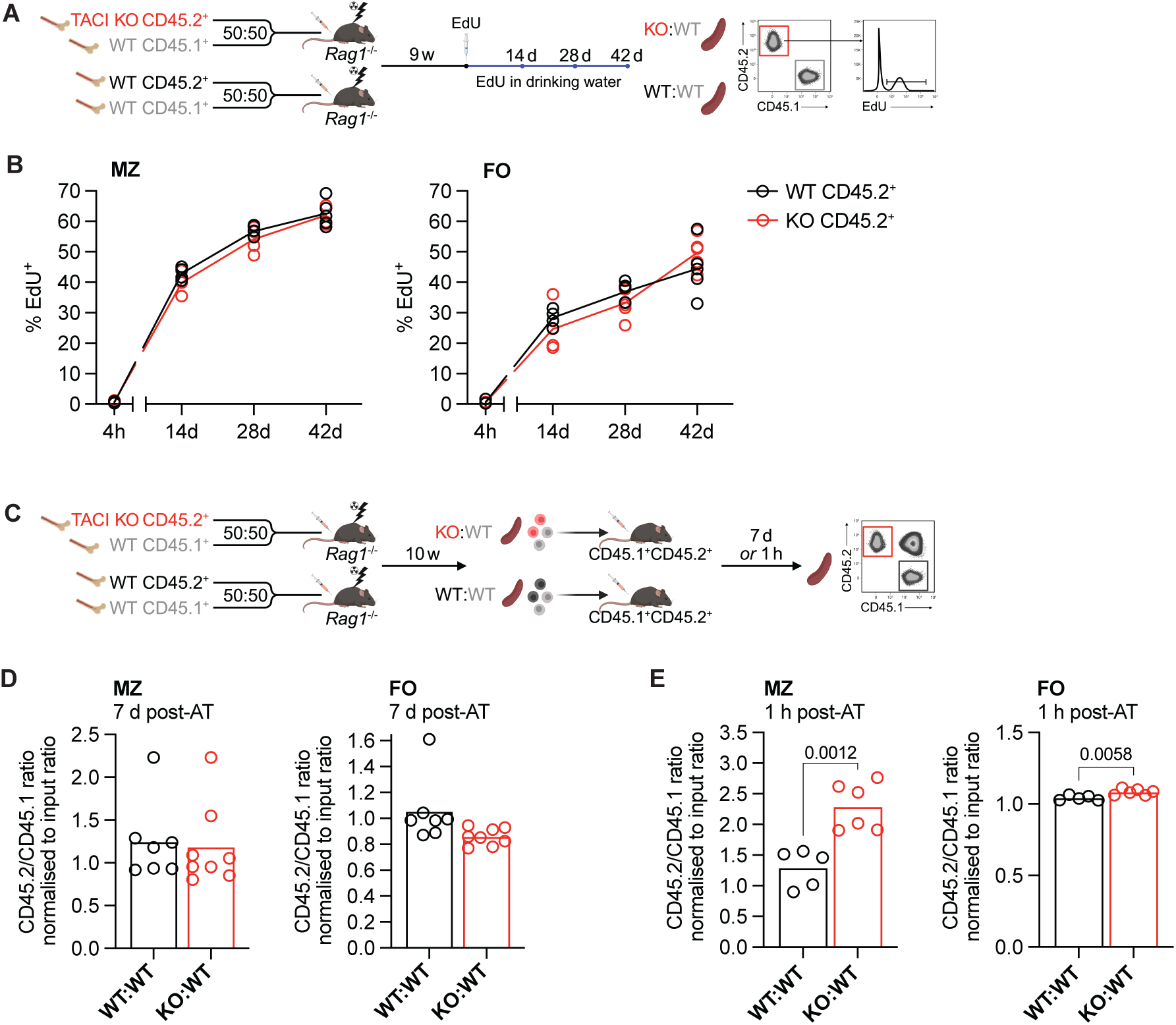
TACI-deficient MZ B cells do not have a survival or homing defect. **A.** Schematic of *in vivo* continuous EdU labelling. WT:WT and KO:WT mixed bone marrow chimeras were generated as in Fig. 1B. 9 weeks later mice were injected i.p. once with EdU and then EdU was maintained in drinking water for 42 days. Spleens were harvested at 4 h, and 14, 28 and 42 days following EdU injection. **B.** The proportion of EdU^+^ cells within MZ (CD1d^hi^IgM^hi^) and FO (CD1d^med^IgM^+^) CD45.2^+^ mature splenic B cell populations (CD93^-^B220^+^CD19^+^) from WT:WT and KO:WT chimeras at 4 h, 14 days, 28 days and 42 days after start of EdU treatment. Cells were analyzed by flow cytometry using the gating strategy shown in Fig. S3A. Each point represents one mouse (4 h WT n = 4, KO n = 5; 14 d n = 4; 28 d and 42 d n = 5). Lines connect the mean at each timepoint. Black: WT CD45.2^+^; red: TACI KO CD45.2^+^. **C.** Schematic of adoptive transfer. WT:WT and KO:WT mixed bone marrow chimeras were generated as in Fig. 1B. At least 10 weeks post-reconstitution, splenic B2 cells were harvested from chimeras and transferred by i.p. injection into CD45.1^+^CD45.2^+^ recipients. Spleens of recipient mice were harvested either 7 days or 1 h later. The ratio of CD45.2^+^/CD45.1^+^ cells recovered was normalized to the ratio of CD45.2^+^/CD45.1^+^ cells in the injected mix (input). **D.** The ratio of CD45.2^+^ to CD45.1^+^ MZ B cells (CD1d^hi^IgM^hi^CD93^-^B220^+^CD19^+^) and FO B cells (CD1d^med^IgM^+^CD93^-^B220^+^CD19^+^) recovered from the spleens of recipient mice 7 days after adoptive transfer (AT) of B cells from WT:WT or KO:WT chimeras, normalized to the ratio of CD45.2^+^ to CD45.1^+^ MZ or FO B cells in the B cells that were transferred (input). Cells were analyzed by flow cytometry using the gating strategy shown in Fig. S3C. Bars show the mean of n = 7 (WT:WT) and n = 8 (KO:WT) recipient mice. Each point represents one recipient mouse. **E.** The ratio of CD45.2^+^ to CD45.1^+^ MZ B cells (CD1d^hi^IgM^hi^CD93^-^B220^+^CD19^+^) and FO B cells (CD1d^med^IgM^+^CD93^-^B220^+^CD19^+^) recovered from the spleens of recipient mice 1 hour after adoptive transfer of B cells from WT:WT or KO:WT chimeras, normalized to the ratio of CD45.2^+^ to CD45.1^+^ MZ or FO B cells in the B cells that were transferred (input). Cells were analyzed by flow cytometry using the gating strategy shown in Fig. S3D. Bars show the mean of n = 5 (WT:WT) and n = 6 (KO:WT) recipient mice. Each point represents one recipient mouse. Statistical tests: two-way ANOVA with Šídák’s multiple comparisons test (B), unpaired t test (D, E). Numbers above graphs indicate p-values where p≤0.05 otherwise no value is shown. Data pooled from 1 (B) or 2 independent experiments (D, E).

We next considered whether loss of TACI affected homing of MZ B cells to the spleen. To examine this, we repeated the adoptive transfer experiment but analyzed the recipient mice 1 hour post-transfer. We found that TACI-deficient MZ and FO B cells homed more efficiently to the spleen compared to WT cells (Fig. 3E, S3D). Thus, loss of TACI does not impair homing of MZ B cells to the spleen. Taken together these results demonstrate that loss of TACI does not reduce survival or homing of MZ B cells, and instead suggest that the lower numbers of MZ B cells are most likely due to impaired development of MZ B cells from their immature T2 B cell precursors.

### TACI regulates the transcriptome of MZ B cells

MZ and FO B cells develop from transitional T2 B cells in the spleen. Maturation of MZ B cells involves upregulation of CD1d, IgM and CD21 and downregulation of CD23 and IgD surface expression (Fig. S4A). In contrast, FO B cells decrease CD1d, IgM and CD21 expression, increase IgD, and maintain high CD23 levels. Analysis of the expression of these proteins on TACI-deficient MZ B cells from KO:WT chimeras showed that in comparison to WT MZ B cells, they express less CD1d, IgM and CD21 and more CD23 and IgD (Fig. 4A-B). Thus, TACI- deficient MZ B cells are more FO-like and less MZ-like, although still clearly distinct from WT FO cells, and appear to be less mature MZ B cells.

**Figure 4.**
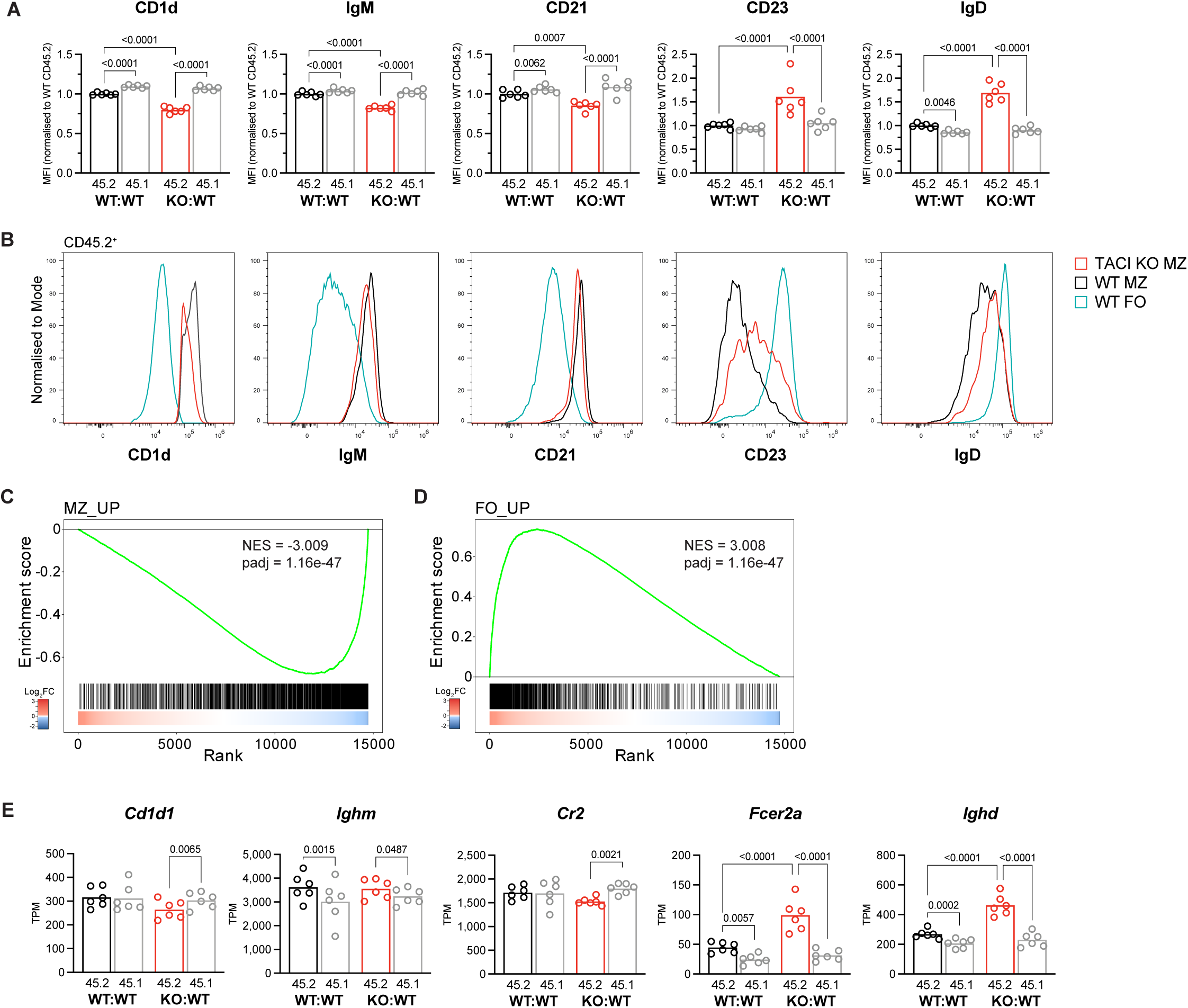
TACI regulates the MZ B cell transcriptome. **A.** CD1d, IgM, CD21, CD23 and IgD surface expression, measured by flow cytometry, on CD45.2^+^ and CD45.1^+^ splenic MZ B cells (CD1d^hi^IgM^hi^CD93^-^B220^+^CD19^+^) from WT:WT and KO:WT chimeras, quantified as the geometric mean fluorescence intensity (MFI) and normalized to the mean MFI of WT CD45.2^+^ cells within the same experiment. Bars show the mean of data pooled from 2 independent analyses. Each point represents one mouse. Black: WT CD45.2^+^; red: TACI KO CD45.2^+^; grey: WT CD45.1^+^. **B.** Flow cytometry histograms comparing CD1d, IgM, CD21, CD23 and IgD surface expression on TACI KO and WT CD45.2^+^ MZ B cells (CD1d^hi^IgM^hi^CD93^-^B220^+^CD19^+^) and WT CD45.2^+^ FO B cells (CD1d^med^IgM^+^CD93^-^B220^+^CD19^+^) from WT:WT and KO:WT chimeras. Histograms are representative of the experiments quantified in A. Red: TACI KO CD45.2^+^ MZ; black: WT CD45.2^+^ MZ; blue: WT CD45.2^+^ FO. **C, D.** Gene set enrichment analysis of the MZ signature (MZ_UP gene set) (C) and the FO signature (FO_UP gene set) (D) in TACI KO vs WT CD45.2^+^ splenic MZ B cells. The plot shows the running enrichment score (green line) above the ranked list of genes, ordered from highest to lowest log2-fold change in TACI KO vs WT CD45.2^+^ MZ B cells (red: upregulated; blue: downregulated; white: no change). The MZ_UP gene set comprises 2,339 genes significantly upregulated ≥1.5-fold in MZ B cells compared to FO B cells and the FO_UP gene set comprises 1,321 genes significantly upregulated ≥1.5-fold in FO B cells compared to MZ B cells (WT CD45.2^+^, from this study, Table S2). NES, normalized enrichment score; padj, adjusted p-value. **E.** *Cd1d*, *Ighm*, *Cr2*, *Fcer2a* and *Ighd* mRNA expression in CD45.2^+^ and CD45.1^+^ splenic MZ B cells (CD1d^hi^IgM^hi^CD93^-^B220^+^CD19^+^) from WT:WT and KO:WT chimeras, determined by RNAseq and quantified as transcripts per million (TPM). Bars show the mean. Each point represents one mouse. Black: WT CD45.2^+^; red: TACI KO CD45.2^+^; grey: WT CD45.1^+^. Statistical tests: repeated measures two-way ANOVA with Fisher’s LSD test. Numbers above graphs indicate p-values where p≤0.05 otherwise no value is shown. Sample numbers: 6 (A, E).

To extend this analysis, we used RNA sequencing (RNAseq) to analyze the transcriptomes of TACI-deficient and WT MZ and FO B cells in KO:WT and WT:WT chimeras, including both the CD45.2^+^ and CD45.1^+^ B cells in the analysis (Fig. S4B). Comparison of the transcriptomes of TACI-deficient and WT MZ B cells (CD45.2^+^) revealed 686 differentially expressed genes (DEGs; fold change > 1.2, padj < 0.05) (Fig. S4C, Table S2). In contrast, no genes were differentially expressed when we compared control WT CD45.1^+^ MZ B cells from KO:WT and WT:WT chimeras, and only 34 genes were differentially expressed between TACI-deficient and WT CD45.2^+^ FO B cells (Fig. S4D-E, Table S2). Thus, loss of TACI selectively affects the transcriptome of MZ B cells, and not FO B cells, and the effect is cell-intrinsic.

To investigate whether TACI-deficient MZ B cells had a less MZ-like and more FO-like identity, we generated transcriptional signatures of MZ and FO B cells from a comparison of WT MZ and WT FO B cells (Table S2). Gene set enrichment analysis (GSEA) showed that TACI- deficient MZ B cells had decreased expression of the MZ signature and increased expression of the FO signature (Fig. 4C-D). Furthermore, in agreement with the changes in protein expression at the surface, transcripts encoding CD1d (*Cd1d1*) and CD21 (*Cr2*) were reduced and those encoding CD23 (*Fcer2a*) and IgD (*Ighd*) were increased in TACI-deficient MZ B cells (Fig. 4E). Levels of transcripts encoding IgM (*Ighm*) were unchanged, implying that post-transcriptional mechanisms are likely to be responsible for the altered surface levels of IgM. Taken together these data show that TACI-deficient MZ B cells are less MZ-like than WT MZ B cells and suggest that TACI regulates the development and maturation of MZ B cells from their T2 B cell precursors and the MZ versus FO B cell fate.

### TACI-deficient MZ B cells have increased FOXO1 activity

To gain insights into how TACI may regulate the differentiation of MZ B cells, we investigated the effect of TACI deficiency on transcription factor (TF) activity. Using GSEA to evaluate the expression of 475 TF-target gene sets, we found that FOXO1 and FOXO3 target genes were among the most upregulated in TACI-deficient MZ B cells (Fig. 5A). Additional GSEA using genes that are both bound and activated by FOXO1 in mouse lymphocytes (Table S2) showed increased expression of FOXO1-activated target genes in TACI-deficient MZ B cells (Fig. 5B, Table S2). Furthermore, loss of TACI resulted in increased expression of *Cxcr4*, *Bach2*, *Klf2*, *Ccr7* and *Cd55*, known FOXO1 targets in B and T lymphocytes (Fig. 5C-D) (Chen et al., 2010; Dominguez-Sola et al., 2015; Kim et al., 2013; Ochiai et al., 2012; Ouyang et al., 2012; Spinelli et al., 2021; Tamahara et al., 2017; Webb et al., 2016). In contrast, *Blimp1*, which is repressed by FOXO1 in B cells (Dominguez-Sola et al., 2015), was downregulated in TACI-deficient MZ B cells compared to WT MZ B cells (Fig. 5D). Together, these results indicate that loss of TACI in MZ B cells results in elevated FOXO1 activity.

**Figure 5.**
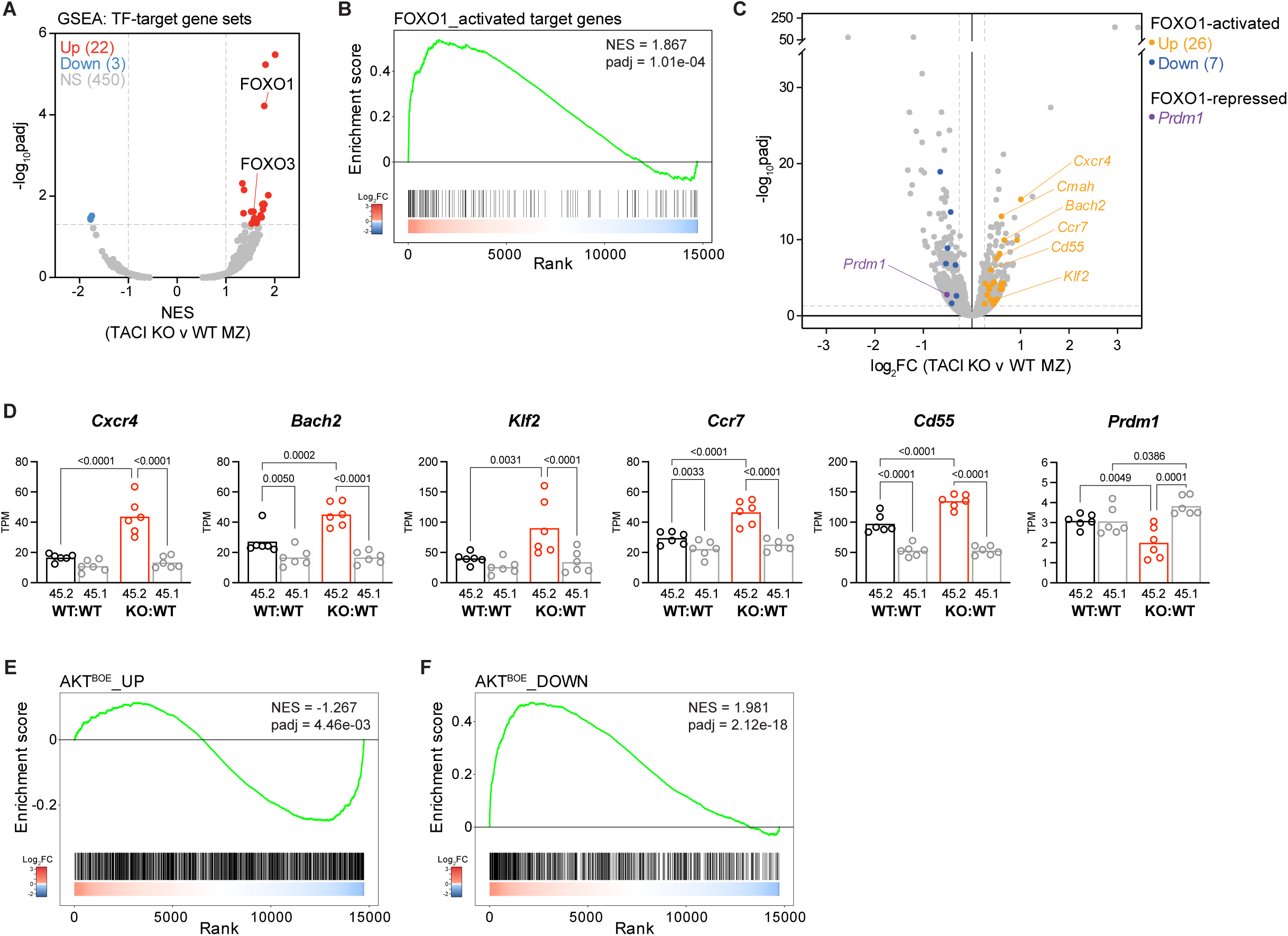
Increased FOXO1 activity in TACI-deficient MZ B cells. **A.** Volcano plot of gene set enrichment in TACI KO vs WT CD45.2^+^ MZ B cells for 475 TF-target gene sets, showing the normalized enrichment score (NES) and -log10-adjusted p-value (-log10padj) for each gene set determined by gene set enrichment analysis (GSEA). Significantly differentially expressed TF-target gene sets are in red (upregulated; NES ≥ 1, padj < 0.05) and blue (downregulated; NES ≤ -1, padj < 0.05). Grey: not significantly differentially expressed (NS). **B.** Gene set enrichment analysis of FOXO1-activated target genes in TACI KO vs WT CD45.2^+^ splenic MZ B cells. The plot shows the running enrichment score (green line) above the ranked list of genes, ordered from highest to lowest log2-fold change in TACI KO vs WT CD45.2^+^ MZ B cells (red: upregulated; blue: downregulated; white: no change). The ‘FOXO1_activated target genes’ gene set comprises 158 genes that are bound by FOXO1 in mouse lymphocytes and are downregulated following FOXO1-knockout, curated as described in Materials & Methods. NES, normalized enrichment score; padj, adjusted p-value. **C.** Volcano plot of differential gene expression between TACI KO and WT CD45.2^+^ MZ B cells from KO:WT and WT:WT chimeras, respectively, with differentially expressed FOXO1- activated or repressed target genes highlighted (orange: FOXO1-activated and upregulated, log2FC ≥ 0.263, padj < 0.05; blue: FOXO1-activated and downregulated, log2FC ≤ -0.263, padj < 0.05; purple: FOXO1-repressed and downregulated). The mean log2-fold change (log2FC) in KO vs WT cells and -log10-adjusted p-value (-log10padj) is shown for each gene from n = 6 mice. **D.** *Cxcr4*, *Bach2*, *Klf2*, *Ccr7*, *Cd55* and *Prdm1* mRNA expression in CD45.2^+^ and CD45.1^+^ splenic MZ B cells (CD1d^hi^IgM^hi^CD93^-^B220^+^CD19^+^) from WT:WT and KO:WT chimeras, determined by RNAseq and quantified as transcripts per million (TPM). Bars show the mean of n = 6 mice. Each point represents one mouse. Black: WT CD45.2^+^; red: TACI KO CD45.2^+^; grey: WT CD45.1^+^. P-values are shown for differences with a p value ≤ 0.05, as determined by repeated measures two-way ANOVA with Fisher’s LSD test. **E, F.** Gene set enrichment analysis of AKT-positively regulated genes (AKT^BOE^_UP gene set) (E) and of AKT-negatively regulated genes (AKT^BOE^_DOWN gene set) (F) in TACI KO vs WT CD45.2^+^ splenic MZ B cells, displayed as in B. The AKT^BOE^_UP gene set comprises 1,478 genes that are upregulated and the AKT^BOE^_DOWN gene set comprises 909 genes that are downregulated in B cells overexpressing constitutively activated AKT1 (AKT^BOE^), compared to control B cells. NES, normalized enrichment score; padj, adjusted p-value.

FOXO1 is inactivated by the serine/threonine kinases AKT1 and AKT2 (AKT). Active AKT translocates to the nucleus and phosphorylates FOXO1, leading to FOXO1 nuclear exclusion and cytosolic sequestration, thereby reducing the expression of FOXO1 target genes. Conversely, loss of AKT activity causes nuclear localization of FOXO1. Furthermore, it has been shown that AKT-mediated inactivation of FOXO1 is critical for MZ B cell development (Cox et al., 2023). Given the increased FOXO1 activity in TACI-deficient MZ B cells, we considered whether this may be caused by reduced AKT activation. We performed GSEA using sets of genes that are up- or down-regulated in B cells following expression of constitutively active AKT1 (Table S2) (Cox et al., 2023). This showed that most of the genes upregulated by AKT1 overactivation were downregulated in TACI-deficient MZ B cells, whereas conversely, genes downregulated by AKT1 overactivation were increased in expression (Fig. 5E-F). These analyses indicate that loss of TACI results in decreased AKT activity and increased FOXO1 activity in MZ B cells, suggesting that TACI regulates the AKT- FOXO1 pathway required for MZ B cell development.

### AKT-mTORC1 signaling is impaired in TACI KO MZ B cells

Given the indication of reduced AKT1 activity in the absence of TACI, we examined phosphorylation of AKT (pAKT) at two sites required for its full activation, T308 and S473, in MZ B cells from KO:WT and WT:WT chimeras. We found that TACI-deficient MZ B cells had reduced resting levels of pAKT at both T308 and S473, compared to WT CD45.2^+^ MZ B cells and to internal control WT CD45.1^+^ MZ B cells, in line with reduced AKT activation (Fig. 6A- D).

**Figure 6.**
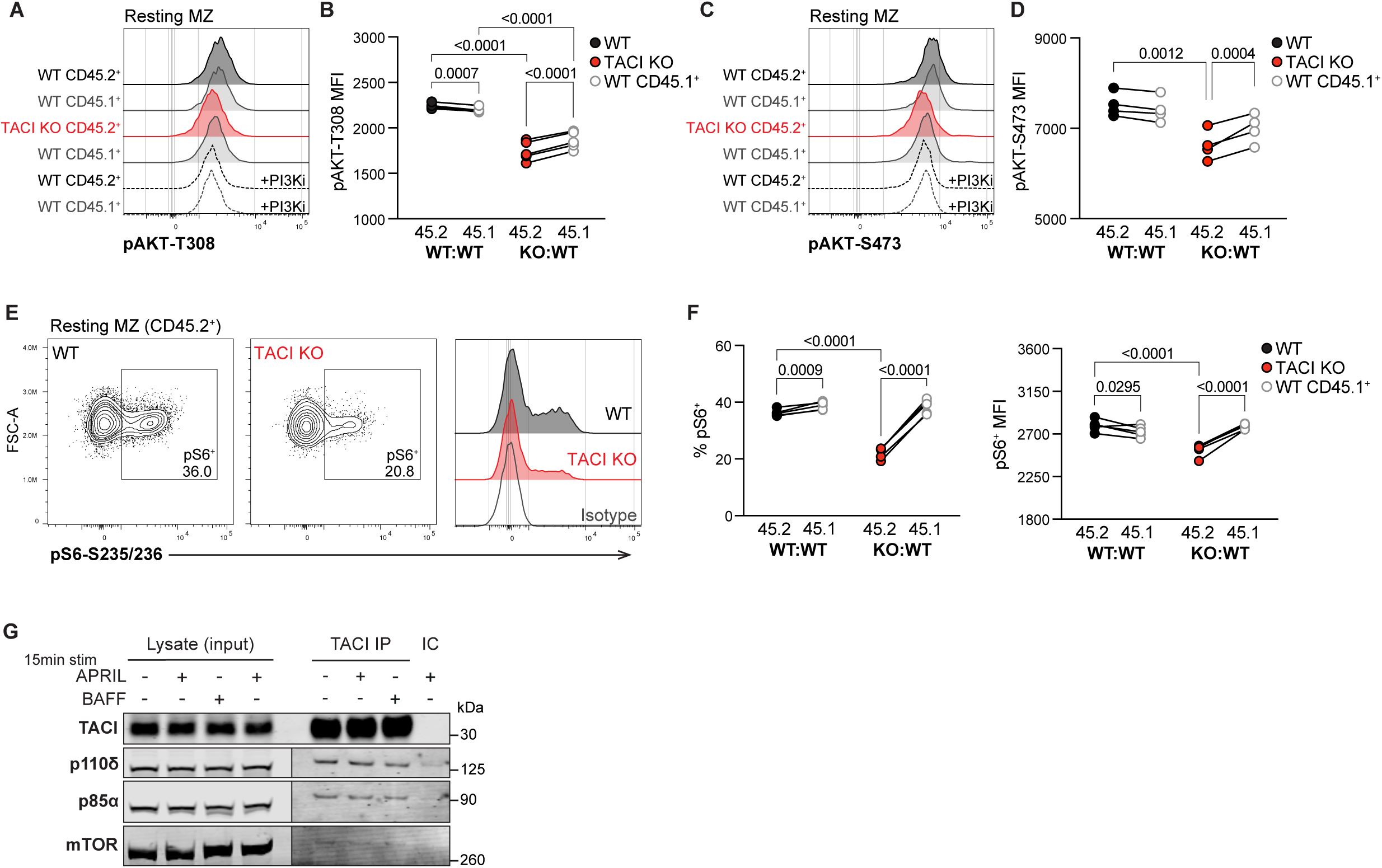
AKT-mTORC1 signaling is impaired in TACI KO MZ B cells. **A.** Flow cytometry histograms comparing pAKT-T308 levels in TACI KO CD45.2^+^, WT CD45.2^+^ and WT CD45.1^+^ resting MZ B cells at 37°C from WT:WT and KO:WT chimeras. Red: TACI KO CD45.2^+^ MZ; black: WT CD45.2^+^ MZ; grey: WT CD45.1^+^; dashed lines: cells treated with the PI3Kδ inhibitor idelalisib (PI3Ki). **B.** pAKT-T308 MFI, measured by flow cytometry, in CD45.2^+^ and CD45.1^+^ resting MZ B cells from WT:WT and KO:WT chimeras. WT:WT n = 4 and KO:WT n = 5, from one of four independent experiments. Lines connect cells from the same sample. Black: WT CD45.2^+^; red: TACI KO CD45.2^+^; grey: WT CD45.1^+^. Numbers above graphs indicate p-values where p≤0.05, as determined by repeated measures two-way ANOVA with Fisher’s LSD test. **C.** Flow cytometry histograms comparing pAKT-S473 levels in TACI KO CD45.2^+^, WT CD45.2^+^ and WT CD45.1^+^ resting MZ B cells at 37°C from WT:WT and KO:WT chimeras. Red: TACI KO CD45.2^+^ MZ; black: WT CD45.2^+^ MZ; grey: WT CD45.1^+^; dashed lines: cells treated with the PI3Kδ inhibitor idelalisib (PI3Ki). **D.** pAKT-S473 MFI, measured by flow cytometry, in CD45.2^+^ and CD45.1^+^ resting MZ B cells from WT:WT and KO:WT chimeras. n = 4, from one of two independent experiments. Lines connect cells from the same sample. Black: WT CD45.2^+^; red: TACI KO CD45.2^+^; grey: WT CD45.1^+^. Numbers above graphs indicate p-values where p≤0.05 otherwise no value is shown, as determined by repeated measures two-way ANOVA with Fisher’s LSD test. **E.** Flow cytometry plots and histograms comparing pS6-S235/236 levels in WT and TACI KO CD45.2^+^ resting MZ B cells at 37°C from WT:WT and KO:WT chimeras. Black: WT CD45.2^+^; red: TACI KO CD45.2^+^; grey: WT cells stained with isotype control antibody (Isotype). **F.** The percentage of pS6^+^ cells (left) and pS6 MFI within pS6^+^ cells (right), measured by flow cytometry, in CD45.2^+^ and CD45.1^+^ resting MZ B cells from WT:WT and KO:WT chimeras. N = 5, from one of four independent experiments. Lines connect cells from the same sample. Black: WT CD45.2^+^; red: TACI KO CD45.2^+^; grey: WT CD45.1^+^. Numbers above graphs indicate p-values where p≤0.05 otherwise no value is shown, as determined by repeated measures two-way ANOVA with Fisher’s LSD test. **G.** Immunoblot anti-TACI immunoprecipitation (IP) from LPS-activated B cells, alongside a control IP using isotype-control antibody (IC) and the input cell lysates. Cells were stimulated with APRIL, BAFF or media-only control for 15 min at 37°C. The immunoblot was probed with anti-TACI, anti-p110δ, anti-p85α and anti-mTOR. The same membrane sections are displayed at two different image intensities for p110δ, p85α and mTOR due to higher protein levels in the input lysates. Representative of two independent experiments.

Besides FOXO1, a key effector of AKT is the mTOR complex 1 (mTORC1). AKT phosphorylates and inhibits TSC1 and TSC2, negative regulators of mTORC1, thereby promoting mTORC1 activity. Activated mTORC1 phosphorylates and activates S6K1, which in turn phosphorylates S6 ribosomal protein at S235 and S236 (pS6). Therefore, we examined whether mTORC1 signaling was impaired in TACI-deficient MZ B cells by measuring pS6, which is mTORC1-dependent in MZ B cells (Fig. S4F). We found that TACI-deficient MZ B cells had reduced levels of pS6 compared to WT MZ B cells (Fig. 6E-F). Thus, loss of TACI in MZ B cells results in reduced AKT and mTORC1 activity, suggesting that TACI regulates both kinases.

### TACI associates with PI3K and mTOR

AKT activation is regulated by PI3K activity, and mice expressing kinase-inactive PI3Kδ, the dominant Class IA PI3K isoform in lymphocytes, lack MZ B cells (Okkenhaug et al., 2002). PI3Kδ activation by cell-surface receptors necessitates its recruitment to the plasma membrane where it generates phosphatidylinositol (3,4,5)-trisphosphate (PIP3), which in turn recruits AKT to enable its phosphorylation and activation by PDK1. Thus, we hypothesized that TACI may regulate membrane localization of PI3Kδ. In agreement with this, we found that TACI associated with both the p85α regulatory subunit and the p110δ catalytic subunit of PI3Kδ (Fig. 6G). Notably, TACI also associated with mTOR (Fig. 6G). All three associations were seen both in the absence and presence of BAFF or APRIL and thus are independent of ligand engagement by TACI. These results suggest that TACI associates with PI3K and mTOR to regulate AKT and mTORC1 signaling.

## Discussion

Our study of the BAFFR interactome revealed that TACI associates with BAFFR. Since the BAFFR-TACI interaction was dependent on the presence of BAFF, the interaction may be ligand-mediated. Both BAFFR and TACI form homotrimers that can bind trimeric BAFF. However, BAFF trimers can interact with each other to form higher order oligomers, a process that is essential for BAFF signaling through BAFFR (Vigolo et al., 2018). We speculate that such BAFF oligomers could simultaneously bind BAFFR and TACI trimers, thus bridging their interaction and promoting signaling through both receptors. Alternatively, BAFFR and TACI could interact directly. The functional consequence of the BAFFR-TACI interaction is unclear; it does not appear to be required for the survival of either FO or MZ B cells, as these are both unaffected by loss of TACI, but it may be required for effective signaling through TACI to regulate MZ B cell development.

Previous studies had shown that loss of TACI leads to an increased number of both FO and MZ B cells, leading to the conclusion that TACI is a negative regulator of B cell survival (Shulga-Morskaya et al., 2004; von Bulow et al., 2001; Yan et al., 2001). However, since TACI- deficient mice have elevated levels of BAFF in the serum (Bossen et al., 2008; Sintes et al., 2017), and overexpression of BAFF in mice leads to more B cells (Mackay et al., 1999), the increased number of B cells may be a direct consequence of higher serum BAFF. Our results with mixed chimeras support this conclusion: the KO:WT chimeras no longer had elevated levels of BAFF, B cell hyperplasia, or increased ICOSL expression, a hallmark of elevated BAFF-BAFFR signaling (Ou et al., 2012). The increase in serum BAFF in TACI-deficient mice may reflect a function of TACI as a decoy receptor for BAFF; ADAM10-mediated cleavage of TACI releases its extracellular domain which can bind and sequester BAFF (Hoffmann et al., 2015; Smulski and Eibel, 2018). Alternatively, TACI may bind and internalize BAFF and target it for degradation, thereby contributing indirectly to B cell homoeostasis.

The analysis of mixed chimeras eliminated the elevated BAFF, B cell hyperplasia and increased BAFFR signaling seen in TACI-deficient mice, allowing us to reveal a cell-intrinsic role for TACI. We show that loss of TACI leads to fewer MZ B cells because of impaired development of the cells from T2 B cell precursors. Surprisingly, TACI was not required for the survival of MZ or FO B cells. In line with this developmental role for TACI, expression of the receptor is low on both T1 and T2 B cells and is upregulated strongly as they mature, especially in MZ B cells. Our results are supported by a previous study of mice expressing a

TACI-Ig fusion protein which binds and sequesters both BAFF and APRIL (Tardivel et al., 2004). These mice have very few FO or MZ B cells, as expected given the essential role of BAFF-BAFFR signaling in mature B cell survival. However, ectopic expression of the anti-apoptotic protein BCL2 in these mice rescued FO B cell numbers, but not MZ numbers, indicating that BAFF and/or APRIL provide a survival-independent signal required for the maturation of MZ B cells. We propose that this signal is mediated by TACI, though there may also be a contribution from BAFFR. In BAFFR-deficient and A/WySnJ (BAFFR-mutated) mice, which have very few mature B cells, forced expression of BCL2 again restores numbers of FO but not MZ B cells, suggesting that BAFFR may also play a role in MZ development (Rahman and Manser, 2004; Sasaki et al., 2004). With respect to whether BAFF, APRIL, or both signal via TACI for MZ B cell development, we note that APRIL-deficient mice have normal numbers of MZ B cells (Varfolomeev et al., 2004). Similarly, mice expressing a BCMA-Ig fusion protein that binds and sequesters APRIL also have unaffected numbers of FO and MZ B cells (Schneider et al., 2001). BCMA is a third member of the TNF receptor subfamily that includes BAFFR and TACI, and binds APRIL but not BAFF. Taken together, these studies demonstrate that BAFF alone is sufficient to support TACI-dependent MZ B cell development. Further studies to support this conclusion would require the generation of mice with mutations that selectively affect BAFF binding to TACI without affecting the BAFF-BAFFR interaction critical for survival.

Our analysis of TACI-dependent signaling and gene expression in MZ B cells showed that loss of TACI results in reduced activation of AKT and increased FOXO1 activity. Given that AKT phosphorylates FOXO1 leading to its nuclear exclusion, the activation of FOXO1 in TACI- deficient MZ B cells is most likely due to reduced AKT activity, which will lead to increased active FOXO1 in the nucleus. The activation of AKT requires PI3K activity, and we show that TACI constitutively associates with the p85α regulatory subunit and p110δ catalytic subunit of PI3Kδ. It is unclear how this interaction is mediated, but it may involve the adaptor MyD88, which has been shown to interact with TACI and can also bind to BCAP, an adaptor protein that in turn binds p85 regulatory subunits of PI3K (He et al., 2010; Okada et al., 2000; Troutman et al., 2012). Given that the BCAP-p85 interaction requires tyrosine phosphorylation of BCAP, the formation of a TACI-MyD88-BCAP-PI3K complex would require kinase activity. This could potentially be provided by BCR signaling leading to activation of the SYK kinase. Interestingly, it has been shown that APRIL induces PI3K-AKT-mTOR signaling in human B cells, which is thought to occur via TACI given that these cells do not express BCMA (Sintes et al., 2017). Thus, we propose that TACI transduces signals via PI3K, localizing the enzyme at the plasma membrane where it can act on its substrate phosphatidylinositol (4,5)- bisphosphate (PIP2) to generate PIP3, leading to AKT recruitment, activation, and subsequent FOXO1 inhibition.

Numerous previous studies support the importance of the PI3K-AKT-FOXO1 pathway in MZ B cell development. Mice expressing a kinase-inactive PI3Kδ lack MZ B cells (Okkenhaug et al., 2002) and deletion of both *Akt1* and *Akt2* in B cells results in a defect in MZ B cell development (Calamito et al., 2010). In contrast, mice expressing hyperactive PI3Kδ in B cells have increased MZ B cell numbers (Stark et al., 2018), while in those overexpressing constitutively membrane-localized and activated AKT1 in B cells, around 90% of splenic B cells are MZ B cells (Cox et al., 2023). Expression of a mutated FOXO1 that cannot be phosphorylated by AKT and hence is constitutively localized to the nucleus results in an almost complete lack of MZ B cells (Cox et al., 2023). Conversely, deletion of FOXO1 in B cells results in an increase in MZ B cells (Chen et al., 2010). One target gene through which FOXO1 may impact on MZ development is *Klf2*. FOXO1 directly binds to the *Klf2* gene and FOXO1- deficient B cells have reduced *Klf2* expression (Chen et al., 2010; Webb et al., 2016). Importantly, mice with KLF2-deficient B cells have more MZ B cells and the remaining FO B cells are more MZ-like, implying that KLF2 may suppress MZ identity (Hart et al., 2011). In alignment with this, we showed that loss of TACI results in increased *Klf2* expression, which may in part account for the reduced numbers of MZ B cells, and for the remaining MZ B cells being less MZ-like. Taken together, these observations support our hypothesis that TACI-mediated activation of PI3K-AKT signaling and subsequent inhibition of FOXO1, resulting in reduced *Klf2* expression, are required for MZ B cell development from T2 B cells.

Several lines of evidence indicate that TACI signaling also activates mTORC1 in MZ B cells. Human and mouse MZ B cells exhibit higher TACI expression and mTORC1 signaling compared to FO B cells, and APRIL has been shown to induce mTORC1 signaling in human B cells, most likely acting through TACI (Gaudette et al., 2020; Sintes et al., 2017). We now show that TACI-deficient mouse MZ B cells have lower basal mTORC1 signaling, in line with reduced AKT activity. Furthermore, we show that TACI associates with mTOR, an interaction that is likely mediated via MyD88 in human B cells (Sintes et al., 2017). Thus, TACI may promote mTORC1 activation in two ways: firstly, by bringing mTOR to the membrane and secondly, by activating AKT, which subsequently phosphorylates and inhibits the negative regulators of mTORC1, TSC1 and TSC2. mTORC1 signaling may establish a poised state in MZ B cells that enables them to undergo rapid plasma cell differentiation (Gaudette et al., 2021; Sintes et al., 2017). mTORC1 regulates cell metabolism, including protein synthesis, as well as the expression of unfolded protein response genes, which are higher in MZ B cells in concordance with their plasma cell-primed state (Gaudette et al., 2020). Furthermore, mTOR-deficient mouse B cells show impaired class-switch recombination and plasmablast differentiation *in vitro* in response to APRIL (Sintes et al., 2017). Our results support the notion that TACI contributes to the unique poised state of MZ B cells by promoting mTORC1 signaling. This, along with the reduced numbers of MZ B cells, may account for the impaired plasma cell expansion in response to a T-independent immunization.

Besides TACI, several other receptors also play important roles in MZ B cell development. BCR signaling is required for both FO and MZ B cell maturation, with MZ B cell development being dependent on BCRs with sufficient auto-reactivity in order to meet the higher threshold of BCR signaling required to enter this compartment, compared to the FO compartment (Noviski et al., 2019). Deletion of the BCR-coreceptor CD19, or mutation of its intracellular domain to impair PI3K binding and activation, causes a near complete loss of MZ B cells (Chen et al., 2010; Martin and Kearney, 2000; Wang et al., 2002). The NOTCH2 receptor also regulates the MZ fate, as loss of NOTCH2 in B cells results in a reduction of MZ B cells, whereas ectopic expression of the NOTCH2 intracellular domain (ICD) causes a large expansion in MZ B cell numbers, in part because of transdifferentiation of FO B cells into MZ B cells (Hampel et al., 2011; Lechner et al., 2021; Saito et al., 2003). Notably, expression of the NOTCH2 ICD results in reduced expression of *Foxo1* and *Klf2* (Lechner et al., 2021). Given that the BCR, CD19 and NOTCH2 activate PI3K and AKT (Aiba et al., 2008; Bailis and Pear, 2012; Castello et al., 2013; Gentle et al., 2012; Otero et al., 2001), these receptors may act in concert with TACI, converging on the same pathways that regulate MZ B cell development. Further work will be needed to understand how signals from these three receptors intersect and their relative contributions to MZ B cell development.

In summary, our results uncover a previously hidden role for TACI in MZ B cell development from T2 B cells. We show that loss of TACI reduces MZ B cell numbers, impairs the T-independent antibody response, and dysregulates the transcriptional programs distinguishing MZ and FO B cells. We propose that TACI regulates MZ development through promoting the PI3K-AKT pathway leading to inhibition of FOXO1 and reduced *Klf2* expression. Our work positions TACI among key receptors, alongside the BCR and NOTCH2, which govern the cell-fate decision between FO and MZ B cells, and broadens our view of the roles played by members of the TNF-receptor superfamily in the establishment and maintenance of the mature B cell repertoire.

## Materials and Methods

### Mice

Mice with the following alleles have been described previously: *Tnfrsf13b*^tm1Vmd^ (*Tnfrsf13b*^-^, TACI KO), *Rag1*^tm1Mom^ (*Rag1*^-^) (Mombaerts et al., 1992; Yan et al., 2001). Mice expressing a tagged BAFFR-TwinStrepTag (*Tnfrsf13c*^em1Tyb^, *Tnfrsf13c*^TwinStrepTag^) were generated using CRISPR/Cas9 gene editing to introduce a mutation that adds 32 amino acids to the C-terminus of BAFFR, encoding a linker and two StrepTag II sequences separated by another linker (amino acid sequence: 172-GPEQ<u>GGGSWSHPQFEKGGGSGGGSGGGSWSHPQFEK</u>*-208; DNA sequence: 5’-GGC CCA GAG CAA <u>GGT GGA GGA TCT TGG AGC CAC CCT CAATTC GAG AAG GGT GGA GGC AGT GGC GGG GGT TCT GGC GGC GGT TCA TGG TCCCAC CCC CAG TTT GAA AAA TAG CCT TGC CGA GGC CAT GG</u>C AGC AGT GGA-3’; underlined is inserted, non-underlined is wildtype) (Fig. S1A). TACI KO mice were on a C57BL/6NCrl background and maintained by intercrossing heterozygous mutants, taking homozygous mutant and wildtype littermate mice for experiments. RAG1 KO and BAFFR- TwinStrepTag mouse strains were backcrossed ≥ 10 generations onto the C57BL/6J (B6) background. B6 (CD45.2^+^Ly9.1^-^), B6.SJL-*Ptprc^a^Pepc^b^*/BoyJ (CD45.1^+^) and 129S7 (129, CD45.2^+^Ly9.1^+^) strains were maintained as inbred strains by the Francis Crick Institute Biological Research Facility, who also generated (B6 x B6.SJL)F1 (CD45.1^+^CD45.2^+^) hybrid mice.

Mice were housed in specific pathogen-free conditions with up to 5 mice per individually ventilated cage (floor area 500 cm^2^; Tecniplast) with bedding, nesting and enrichment (Aspen 4HK, Bed-R’Nest; Datesand) at 21 ± 2°C and 55 ± 10% relative humidity in a 12 h/12 h light/dark cycle. Mice were fed 2018 Teklad Global Diet (Envigo; autoclaved) and supplied RO-filtered mains drinking water, *ad libitum*. Both female and male mice were used, with sexes balanced between different genotypes or conditions. All procedures were conducted in accordance with the United Kingdom Animal (Scientific Procedures) Act 1986, approved by the Francis Crick Institute Animal Welfare and Ethical Review Body and conducted under authority of a Project Licence issued by the UK Home Office.

### Mixed bone marrow chimeras

Bone marrow was isolated from femurs and tibias in IMDM media (in-house, with 0.06 mg/ml penicillin, 0.1 mg/ml streptomycin, pH 7.2) supplemented with 5% FCS, then red blood cells were lysed with ACK Lysing Buffer (Gibco) for 2 min at room temperature. T cells were depleted by incubation with biotinylated anti-CD3ε (100304, BioLegend; 2.5 μg/ml) for 20 min on ice and then with streptavidin-conjugated Dynabeads (M-280, Thermo Fisher Scientific; 1 × 10^6^ beads per 1 × 10^6^ cells) for 20 min at 4°C with rotation, followed by removal of the bead-bound cells with a magnet, washing and resuspension in IMDM without FCS. T-cell depleted bone marrow from *Tnfrsf13b*^+/+^ (WT) or *Tnfrsf13b*^-/-^ (TACI KO) littermates was mixed 50:50 with bone marrow from either 2-3 B6.SJL (CD45.1^+^) mice, for B6:B6.SJL (CD45.2^+^:CD45.1^+^) chimeras, or one 129S7 mouse, for B6:129 (Ly9.1^-^:Ly9.1^+^) chimeras. 100 μl of bone marrow mix at 2 × 10^7^ cells/ml was then injected into the tail vein of each *Rag1*^-/-^ recipient, which had been irradiated with a sub-lethal dose of 5 Gy from a ^137^Cs source at least 3 h earlier. Donor and recipient mice were sex-matched and aged 8-14 weeks or 7-10 weeks, respectively. Recipients received 0.02% enrofloxacin (Baytril, Bayer healthcare) in their drinking water for 4 weeks and mixed chimeras were analyzed at least 8 weeks post-bone marrow injection to allow the hematopoietic compartment to reconstitute.

### Flow Cytometry and cell sorting

Spleens were harvested and homogenized through 40 μm cell strainers with ice-cold FACS buffer (PBS [Ca^2+^/Mg^2+^-free], 0.5% BSA [A2153, Sigma], 2 mM EDTA) then red blood cells were lysed with ACK Lysing Buffer for 2 min at room temperature. For flow cytometric analyses of splenocytes from chimeras, unless otherwise specified, cells were stained with viability dye diluted 1:500 in PBS at 2 × 10^7^ cells/ml for 20 min at 4°C, then incubated with fluorophore-conjugated antibodies specific for surface markers diluted 1:200 in FACS buffer at 2 × 10^7^ cells/ml for 30 min at 4°C, fixed in 100 μl Cytofix/Cytoperm (554722, BD Biosciences) for 15 min on ice, and analyzed in FACS buffer containing a set number of Calibrite beads (664912, BD Biosciences) on an Aurora spectral flow cytometer (Cytek). All antibody incubations included anti-CD16/32 (101302, BioLegend) to block Fc-receptor binding. Fluorophore-conjugated reagents used were as follows: B220-PerCP-Cy5.5 (RA3-6B2), B220-BV785 (RA3-6B2), B220-BV650 (RA3-6B2), CD138-BV785 (281-2), CD19-BV785 (6D5), CD19-APC (6D5), CD1d-PE (1B1), CD1d-BV421 (1B1), CD23-BV605 (B3B4), CD23-PE-Cy7 (B3B4), CD45.1-BV605 (A20), CD45.2-BV650 (104), CD45.2-BV785 (104), CD45.2-PerCP-Cy5.5 (104), CD93-PE-Cy7 (AA4.1), IgD-BV711 (11-26c.2a), IgD-AF647 (11-26c.2a), IgM-BV510 (RMM-1), IgM[a]-PerCP-Cy5.5 (MA-69), IgM[b]-PE (AF6-78), Streptavidin-PE-Cy7, TCRβ-

AF700 (H57-597) and Zombie Aqua Fixable Viability Dye, all from BioLegend. B220-RB780 (RA3-6B2), CD19-BUV395 (1D3), CD19-APCeF780 (1D3), CD1d-BV605 (1B1), CD2-BV711 (RM2-5), TACI-BV421 (8F10), CD3-BUV395 (2C11), CD93-BUV737 (AA4.1), IgD-BV786 (11- 26c.2a), IgD[a]-FITC (AMS9.1), IgG1[b]-biotin (B68-2), IgG1[a]-BUV805 (109), IgM-BUV563 (II/41), IgM-BV605 (II/41), Ly9.1-BUV563 (30C7) and TCRβ-BUV395 (H57-597) all from BD Biosciences. B220-APC (RA3-6B2), BAFFR-FITC (eBio7H22-E16), CD21-APC-eF780 (8D9), CD45.1-PerCP-Cy5.5 (A20), CD45.2-FITC (104), CD93-APC (AA4.1), ICOSL-eF660 (B7-H2),

TCRβ-APC-eF780 (H57-597), DAPI (used at 0.1 μg/ml with 10 min incubation), and Live/dead Fixable Near-IR Dead Cell Stain Kit (L10119) from Thermo Fisher Scientific. IgM-APC-ef780 (II/41) from eBioscience.

For isolation of B cell populations by fluorescence-activated cell sorting (FACS) for RNAseq, B2 cells were first isolated from splenocyte single-cell suspensions by negative depletion using anti-CD43 MicroBeads (130-049-801, Miltenyi Biotec) and LS magnetic columns (Miltenyi Biotec), according to the manufacturer’s protocol. B cells were stained with viability dye and fluorophore-conjugated antibodies, then sorted live using a FACSAria III or Fusion (BD Biosciences) into cold RPMI-1640 media (R8758; Sigma-Aldrich) supplemented with 10% heat-inactivated FCS, 100 U/ml penicillin, 100 μg/mL streptomycin, 2 mM L-glutamine (Sigma-Aldrich), and 50 μM β-mercaptoethanol (Thermo Fisher Scientific).

### TNP-Ficoll Immunization

At 9 weeks post-reconstitution, B6:129 (WT:wt and KO:wt) chimeras were injected intraperitoneally with 50 μg of TNP-Ficoll (F-1300, LGC Biosearch Technologies) in 200 μl PBS, or PBS only. At day 7, spleens were harvested, and blood was collected by cardiac puncture into serum gel CAT tubes (41.1378.005, Sarstedt) and allowed to clot at room temperature for 1-2 h. Serum was isolated by centrifugation at 10,000 × g for 10 min at room temperature. Splenocytes were incubated with antibodies against cell surface markers and viability dye for 30 min at 4°C, fixed with 250 μl Cytofix Buffer (554655, BD Biosciences) for 20 min at 4°C, and analyzed in FACS buffer containing a set number of Calibrite beads on a Fortessa Symphony A5 (BD Biosciences).

### ELISA

For the analysis of TNP-specific antibodies in serum by ELISA, 96-well MaxiSorp flat-bottom plates (439454, Thermo Fisher Scientific) were coated with 0.5 μg TNP-BSA (T-5050, LGC Biosearch Technologies) in 100 μl PBS overnight at 4°C followed by 1 h at room temperature and then washed five times with PBST (PBS, 0.01% Tween 20). Wells were blocked with 100 μl 3% BSA in PBS for 2 h at room temperature then washed twice with PBST before addition of serially diluted serum in 3% BSA in PBS and incubation for 2 h at room temperature. Wells were washed five times before addition of 0.1 μg biotinylated anti-IgM[a] (553515, BD Biosciences), anti-IgM[b] (406204, BioLegend), anti-IgG1[a] (553500, BD Biosciences), or anti-IgG1[b] (553533, BD Biosciences), in 100 μl PBST and incubation for 2 h at room temperature. Wells were washed five times before addition of 0.2 μg streptavidin-HRP (SA- 5004-1, Vector Laboratories) in 100 μl PBST and incubation for 2 h at room temperature. Wells were washed five times then incubated with 70 μl TMB substrate (00-4201, Thermo Fisher Scientific) until color developed and the reaction was stopped with 70 μl 2N H2SO4.

Absorbance at 450 nm was measured using a Spark microplate reader (Tecan). Antibody titer was recorded as the absorbance at a dilution in the linear section of the titration curve (1:1600 for IgM^a^, 1:200 for IgM^b^, 1:400 for IgG1^a^, and 1:50 for IgG1^b^).

The concentration of BAFF in serum was measured by ELISA using the Mouse BAFF ELISA Kit (ab119580, Abcam) according to the manufacturer’s instructions, with serially diluted serum incubated in anti-BAFF-coated plates overnight at 4°C and washes performed with PBST. Absorbance at 450 nm was measured using a Spark microplate reader and BAFF concentration was calculated using a standard curve generated with recombinant BAFF.

### EdU labelling *in vivo*

At 9 or 12 weeks post-reconstitution, WT:WT and KO:WT chimeras were injected once intraperitoneally with 100 μl PBS containing 1 mg/ml 5-ethynyl-2’-deoxyuridine (EdU; F463473, Fluorochem), then received 0.3 mg/ml EdU in their drinking water for the duration of the time course, with EdU replenished every 3 days. Spleens were harvested at 4 hours, 14 days, 28 days, and 48 days post-injection, to analyze FO and MZ cells, or at 4 hours, 2 days, 3 days and 5 days post-injection to analyze transitional B cells.

To analyze EdU^+^ cells by flow cytometry, spleens were harvested, red blood cells were lysed, and splenocytes were incubated with antibodies against cell surface markers and viability dye for 30 min at 4°C. Cells were washed then fixed with 4% paraformaldehyde for 15 min at 4°C and permeabilized in Perm/Wash Buffer (554723, BD Biosciences) supplemented with 1% BSA for 15 min at room temperature, before addition of click reaction mix to a final concentration of 2 mM CuSO4, 0.3125 μM AZDye488-picolyl-azide, and 10 mM sodium ascorbate (all from Jena Bioscience) in TBS (50 mM Tris, 150 mM NaCl, pH 7.5) and incubation for 30 min in the dark at room temperature. Cells were washed twice then analyzed in Perm/Wash Buffer on a Fortessa Symphony A5.

### Adoptive transfer of B cells

At 10-15 weeks post-reconstitution, spleens were harvested from 2-3 WT:WT and KO:WT chimeras, red blood cells were lysed, and B2 cells were isolated by negative depletion using anti-CD43 MicroBeads and LS magnetic columns, according to the manufacturer’s protocol, and pooled by genotype. For 7 day adoptive transfer experiments, 1 × 10^7^ purified B cells from WT:WT or KO:WT chimeras in 100 μl IMDM media were injected intravenously into each sex-matched CD45.1^+^CD45.2^+^ recipient mouse. For 1 hour adoptive transfer experiments, purified B cells were first incubated with 1 μM CMFDA (C7025, ThermoFisher Scientific) in PBS for 10 min at 37°C, excess dye was quenched with IMDM media containing 10% FCS for 7 min at 37°C, cells were washed, then 1 × 10^7^ CMFDA-labelled B cells from WT:WT or KO:WT chimeras in IMDM media were injected into each sex-matched CD45.1^+^CD45.2^+^ recipient. In each case, a proportion of the transferred cell suspension was analyzed by flow cytometry to determine the input ratio of CD45.2^+^/CD45.1^+^ cells among the FO and MZ B cells injected.

Recipient spleens were harvested 1 hour or 7 days post-adoptive transfer, red blood cells were lysed, and B2 cells were isolated by negative depletion using anti-CD43 MicroBeads and LS magnetic columns, as described before. Cells were incubated with antibodies against cell surface markers and viability dye for 30 min at 4°C, fixed with 250 μl Cytofix Buffer for 20 min at 4°C, and analyzed in FACS buffer containing a set number of Calibrite beads on a Fortessa Symphony A5. The ratio of CD45.2^+^/CD45.1^+^ cells among the FO and MZ B cells recovered was divided by the corresponding ratio of CD45.2^+^/CD45.1^+^ cells in the FO and MZ populations that were transferred.

### RNAseq

Cell pellets were lysed after FACS in Buffer RLT (QIAGEN) with 500 μM β-mercaptoethanol then frozen until processing. Lysates were homogenized using QIAshredder spin columns then RNA was extracted using the RNeasy Mini or Micro kit (QIAGEN) with DNase digestion on a QIAcube Connect, all according to the manufacturer’s instructions. cDNA libraries were generated using the Watchmaker RNA Library Prep Kit with Polaris Depletion of rRNA and globin transcripts according to the manufacturer’s instructions. Libraries were sequenced on the Illumina NovaSeq X, acquiring 25 million 100 bp paired-end reads per sample.

Alignment of paired-end raw reads was performed with the nf-core rnaseq pipeline v3.6 using the star rsem aligner option and the ENSEMBL GRCm39 release 105 transcriptome (https://zenodo.org/records/15631172). Differential gene expression analysis was performed using the DESeq2 R-package, where adjusted p-values (padj) were calculated using the Benjamini-Hochberg method, with a significance threshold of padj < 0.05 (Love et al., 2014). Gene Set Enrichment Analyses were performed using the FGSEA R-package with genes pre-ranked according to log2-foldchange (Korotkevich et al., 2021). GSEA normalized enrichment scores (NES) were computed by permutation testing (1000 permutations) and adjusted p-values were calculated using Benjamini-Hochberg multiple testing correction, with a significance threshold of padj < 0.05. Gene sets used for GSEA were curated as follows: Transcription factor target gene sets were curated using the OmniPath Python package (Turei et al., 2021). FOXO1-activated target genes were manually curated by cross-referencing FOXO1-targets, identified by ChIP-seq in mouse pre-B cells and Tregs (Ochiai et al., 2012; Ouyang et al., 2012; Webb et al., 2016), with genes downregulated in FOXO1 KO vs WT mouse CD8^+^ CTLs (1.3-fold and padj < 0.05 thresholds applied) (Spinelli et al., 2021). AKT^BOE^_UP and AKT^BOE^_DOWN gene sets contain genes upregulated or downregulated in AKT^BOE^ B cells compared to control FO B cells, respectively (Cox et al., 2023). MZ_UP and FO_UP gene sets comprise genes upregulated or downregulated, respectively, in WT MZ compared to WT FO CD45.2^+^ B cells, determined in the current study (1.5-fold change, padj < 0.05). Differential gene expression results and custom gene sets are available in Table S2. RNAseq data has been deposited with GEO (accession number GSE300247; reviewer token: orwdmyswzpwlngr).

### B cell *in vitro* culture

To generate LPS-blasts, spleens were harvested from mice, red blood cells were lysed, and resting B2 cells were isolated by negative depletion using anti-CD43 MicroBeads and LS magnetic columns (Miltenyi Biotec), according to the manufacturer’s protocol, in sterile FACS buffer. Purified B cells were then cultured at 1 × 10^7^ cells per well in 6-well plates (353046, Corning Inc.) with 40 μg/ml LPS (L3129, Sigma-Aldrich) in RPMI-1640 media supplemented with 10% heat-inactivated FCS, 100 U/ml penicillin, 100 μg/mL streptomycin, 2 mM L-glutamine, and 50 μM β-mercaptoethanol for 4 days (for BAFFR affinity purification) or 3 days (for TACI immunoprecipitation) at 37°C, 5% CO2.

### BAFF and APRIL *in vitro* stimulation and cell lysis

Before *in vitro* stimulation of LPS-blasts, cells were washed by centrifugation and rested in 0.5% FCS RPMI media at 2 × 10^7^ cells/ml for 75 min at 37°C. Cells were then stimulated with 200 ng/ml mouse BAFF (8876-BF-010/CF, R&D Systems) or 200 ng/ml mouse APRIL (7907- AP-010/CF, R&D Systems) in 0.5% FCS RPMI for 15 min at 37°C with 350 rpm shaking. Cells were immediately pelleted at 800 × g for 2 min at 4°C then resuspended in ice-cold lysis buffer (1% IGEPAL CA-630, 50 mM HEPES, 150 mM NaCl, 1 mM EDTA, 1 mM EGTA, 25 mM NaF, 10 mM iodoacetamide, 2.5 mM sodium pyrophosphate, 5 mM β-glycerophosphate, 5 mM sodium orthovanadate and 1X Protease Inhibitor Cocktail [ab270055, Abcam]; 80 μl per 1 × 10^7^ cells) and incubated on ice for 10 min. Lysates were cleared by centrifugation at 16,000 × g for 10 min at 4°C.

### BAFFR affinity purification

For each affinity purification, fresh, cleared lysates from 8 × 10^8^ LPS-blasts (*Tnfrsf13c*^TwinStrepTag/TwinStrepTag^ or wildtype) were incubated with 60 μl washed MagStrep type 3 Strep-TactinXT beads (2-4090-002, IBA Lifesciences) for 20 min at 4°C with rotation. Beads were washed three times with lysis buffer at 4°C with rotation then proteins were released by incubation with biotin elution buffer (100 mM biotin, 200 mM Tris-HCl, 300 mM NaCl, 2 mM EDTA, pH 8) for 10 min, twice, with eluted proteins collected using a DynaMag-2 Magnet (Thermo Fisher Scientific) then snap-frozen.

### nLC-MS/MS Analysis

Biotin-eluted affinity purification samples were reduced with 50 mM DTT for 10 min at 95°C then alkylated with 15 mM iodoacetamide for 20 min at room temperature with shaking in the dark. Proteins were precipitated with 8 volumes 100% acetone overnight at -20°C, followed by centrifugation at 20,000 xg for 30 min at 4°C and two washes with ice-cold 80% acetone and centrifugation at 20,000 xg. Air-dried protein pellets were resuspended with sonication in digestion buffer (100 mM HEPES, 1 M guanidine-HCl, pH 8) then incubated with 500 ng LysC for 3 hours at 37°C with shaking followed by overnight digestion with 500 ng trypsin at 37°C with shaking, then quenched with 10% TFA to pH 2-3. Peptide samples were analyzed by nano-liquid chromatography-tandem mass spectrometry (nLC-MS/MS) using an EvoSep One interfaced via a nano-electrospray ion source onto an Orbitrap Fusion Lumos Tribrid Mass Spectrometer (ThermoFisher Scientific). Peptides were separated using the standardized 30 Samples Per Day (30SPD) pre-formed gradient method. Eluted peptides were ionized by applying a 2.2 kV voltage and introduced to the mass spectrometer as gas-phase ions. MS1 scans were acquired in the Orbitrap mass analyzer with a target of 4 ×10^5^ ions or 50 ms maximum injection time, a mass resolution of 60,000 and a scan range of m/z 375–1,500. Ions with charge 2-5 and intensity ≥10,000 from each MS1 scan were isolated in the quadrupole mass analyzer using a 1.2 m/z window and fragmented using higher-energy collisional dissociation with normalized collision energy of 32%. Ions were excluded from reselection for 15 s after fragmentation within a 10 ppm dynamic exclusion window. MS2 spectra were acquired in the Ion Trap mass analyzer at rapid scan rate with a maximum injection time of 300 ms and AGC target of 1 ×10^4^ ions, with the scan range set automatically based on the isolated precursor.

Acquired MS data were processed in MaxQuant version version 2.6.5.0 and spectra were searched against the mouse UniProt reference proteome (UP000000589, downloaded 20/08/2021) using the Andromeda search engine. Trypsin/P was set as the specific protease with a maximum of two missed cleavage sites allowed. Mass tolerance was set to 20 pmm for the first search and 4.5 ppm for the main search, with fragment mass tolerances set to 0.5 Da. Carbamidomethylation of cysteine was set as a fixed modification while oxidation of methionine, diglycine remnant on lysine, and acetylation of the protein N-terminus were set as variable modifications. A maximum of 5 modifications were allowed per peptide, with a minimum peptide length of 7. An FDR of 1% was used for peptide spectrum matches and protein-level identification, and a minimum of one peptide was required for successful protein identification. Razor peptides were used for label-free quantification (LFQ) and intensity based absolute quantification (iBAQ), with a minimum ratio count of 2 required. Quantitative MS data was analyzed in Perseus version 1.6.15.0 (Tyanova and Cox, 2018; Tyanova et al., 2016). LFQ intensity values for proteins, excluding contaminants, that were quantified in at least one sample were log2-transformed, missing values were imputed with a constant below-minimum value, then two-tailed Student’s t-tests were performed to compare the mean abundance of each protein in BAFF-stimulated BAFFR affinity purifications (APs; n = 3) with its mean abundance in control APs (from wildtype cells; n = 3) or unstimulated BAFFR APs (n = 3). Specific interactors were determined as proteins with a ≥ 2-fold-enrichment in BAFFR APs compared to control APs and a p-value ≤ 0.05. Generated p-values were also corrected for multiple hypothesis testing using the permutation-based false discovery rate (FDR) method in Perseus, with a threshold of 0.20 to generate q-values. Full t-test results and q-values can be found in Table S1. Proteomics data have been deposited to the ProteomeXchange Consortium (http://proteomecentral.proteomexchange.org) via the PRIDE partner repository (Perez-Riverol et al., 2019) with the dataset identifier PXD065033 (reviewer token: 0n5MoD7zKvaY)

### Phosphoflow

Spleens were harvested from chimeras, red blood cells were lysed, and B2 cells were isolated by negative depletion using anti-CD43 MicroBeads and LS magnetic columns (Miltenyi Biotec), according to the manufacturer’s protocol. B cells were rested in 0.5% FCS RPMI-1640 media at 4 × 10^6^ cells/ml for 75 min at 37°C, with 200 nM Idelalisib (7631, R&D Systems) for PI3Kδ-inhibited samples or 10 nM rapamycin (553210, Sigma-Aldrich) for mTORC1-inhibited samples. Cells were stained with anti-CD93 for 30 min on ice, washed, then incubated in 0.5% FCS RPMI at 2 × 10^7^ cells/ml for 5 min or 15 min at 37°C with shaking, with Idelalisib or rapamycin for inhibited control samples. Cells were fixed by addition of pre-warmed Cytofix Buffer for 10 min at 37°C. Fixed cells were stained with anti-CD19 and anti-IgM for 30 min at 4°C, permeabilized with pre-chilled Perm Buffer III (558050, BD Biosciences) for 30 min on ice, then stained with anti-TACI, anti-CD45.1, anti-CD45.2, anti-IgD, anti-B220, anti-CD1d and anti-CD21 for 30 min at 4°C. Cells were incubated with anti-p-S6-S235/236 or rabbit IgG isotype control (9865, 8760, Cell Signaling Technology; 0.13 μg/ml) for 30 min at 4°C, or with anti-p-AKT-T308 or rabbit IgG isotype control (48646, 2985, Cell Signaling Technology; 0.25 μg/ml) and anti-p-AKT-S473 or mouse IgG1κ isotype control (562465, 562292, BD Biosciences; 0.12 μg/ml) overnight at 4°C. Samples were analyzed in FACS buffer on an Aurora spectral flow cytometer (Cytek).

### TACI immunoprecipitation

Fresh, cleared lysates from 2 × 10^7^ LPS-blasts were incubated with 2 μg goat polyclonal anti-mouse-TACI (AF1041, R&D Systems) or 2 μg normal goat IgG (AB-108-C, R&D Systems) for 1 h at 4°C with rotation, followed by addition of 20 μl washed protein-G Dynabeads (10003D, Thermo Fisher) and further incubation for 30 min at 4°C with rotation. Beads were washed three times with lysis buffer at 4°C with rotation then proteins were eluted by incubation at 95°C for 5 min in 1X NuPAGE LDS sample buffer (Thermo Fisher) with 40 mM DTT, and beads were removed using a DynaMag-2 Magnet.

### Immunoblotting

Cell lysates were denatured and reduced in 1X NuPAGE LDS Sample Buffer with 40 mM DTT at 70°C for 10 min. Pre-IP lysates (input) equivalent to 1.25 × 10^6^ cells and half of each IP were subjected to SDS-PAGE on NuPAGE 4-12% Bis-Tris Gels (Thermo Fisher Scientific) and proteins transferred onto Immobilon-FL PVDF membrane (Millipore) using the XCell SureLock Mini-Cell Electrophoresis System and XCell II Blot Module (Thermo Fisher Scientific) according to the manufacturer’s instructions. Membranes were blocked with 5% BSA in TBST (TBS, 0.001% Tween 20) for 1 h at room temperature then incubated sequentially with primary antibodies: anti-mTOR (4517, Cell Signaling Technology), anti- p110δ (34050, Cell Signaling Technology), anti-p85α (4257, Cell Signaling Technology), anti-TACI (93005, Cell Signaling Technology), anti-MyD88 (4283, Cell Signaling Technology) in 5% BSA TBST overnight at 4°C. Washed membranes were incubated with goat anti-rabbit IRDye 680RD (925-68071, LI-COR) or goat anti-mouse IRDye 800CW (925-32210, LI-COR) secondary antibodies at 1:10,000 in 5% BSA TBST for 1 h at room temperature. Washed and dried membranes were imaged using an Odyssey CLx Imaging System (LI-COR).

### Statistical Analysis

Statistical significance was assessed using an unpaired t test, two-way ANOVA with Fisher’s LSD test, two-way ANOVA with Šídák’s multiple comparisons test, or an unpaired Welch’s t test corrected for multiple comparisons using the Holm-Šidák method to generate p-values. Statistical analysis of differential gene expression in the RNAseq data was performed using DESeq2, with adjusted p-values calculated using the Benjamini-Hochberg method. GSEA normalized enrichment scores (NES) were computed by permutation testing (1000 permutations) and adjusted p-values were calculated using Benjamini-Hochberg multiple testing correction. Proteomic data was analyzed using a two-tailed Student’s t-test corrected for multiple hypothesis testing using the permutation-based false discovery rate (FDR) method to generate q-values. P-values, adjusted p-values and q-values ≤ 0.05 were considered significant, rejecting the null hypothesis.

## Data and materials availability

RNAseq data has been deposited with GEO (accession number GSE300247; reviewer token: orwdmyswzpwlngr). Proteomics data have been deposited to the ProteomeXchange Consortium with the dataset identifier PXD065033 (reviewer token: 0n5MoD7zKvaY). Other data and materials are available by request to the authors.

## Acknowledgments

We thank Dinis Calado and Edina Schweighoffer for critical reading of the manuscript. We thank the Biological Research Facility, and the Genomics, Flow Cytometry and Proteomics Science Technology Platforms of the Francis Crick Institute. We thank Vishva Dixit (Genentech) for the TACI-deficient mice. VLJT was supported by the Francis Crick Institute (CC 2080) that receives its core funding from Cancer Research UK (CC 2080), the U.K. Medical Research Council (CC 2080), and the Wellcome Trust (CC 2080).

## Author contributions

Conceptualization: DL, VLJT

Methodology: DL, VLJT

Formal analysis: DL, LV, SB

Investigation: DL, LV

Writing - Original Draft: DL, VLJT

Writing - Review & Editing: DL, VLJT Visualization: DL

Supervision: VLJT

Project administration: VLJT Funding acquisition: VLJT

## Supplementary Information

### Supplementary Tables

Table S1

Mass spectrometric analysis of proteins co-purified with BAFFR-TwinStrepTag

Table S2

Analysis of RNAseq of MZ and FO B cells from KO:WT and WT:WT chimeras

### Supplementary Figure Legends

**Figure S1.**
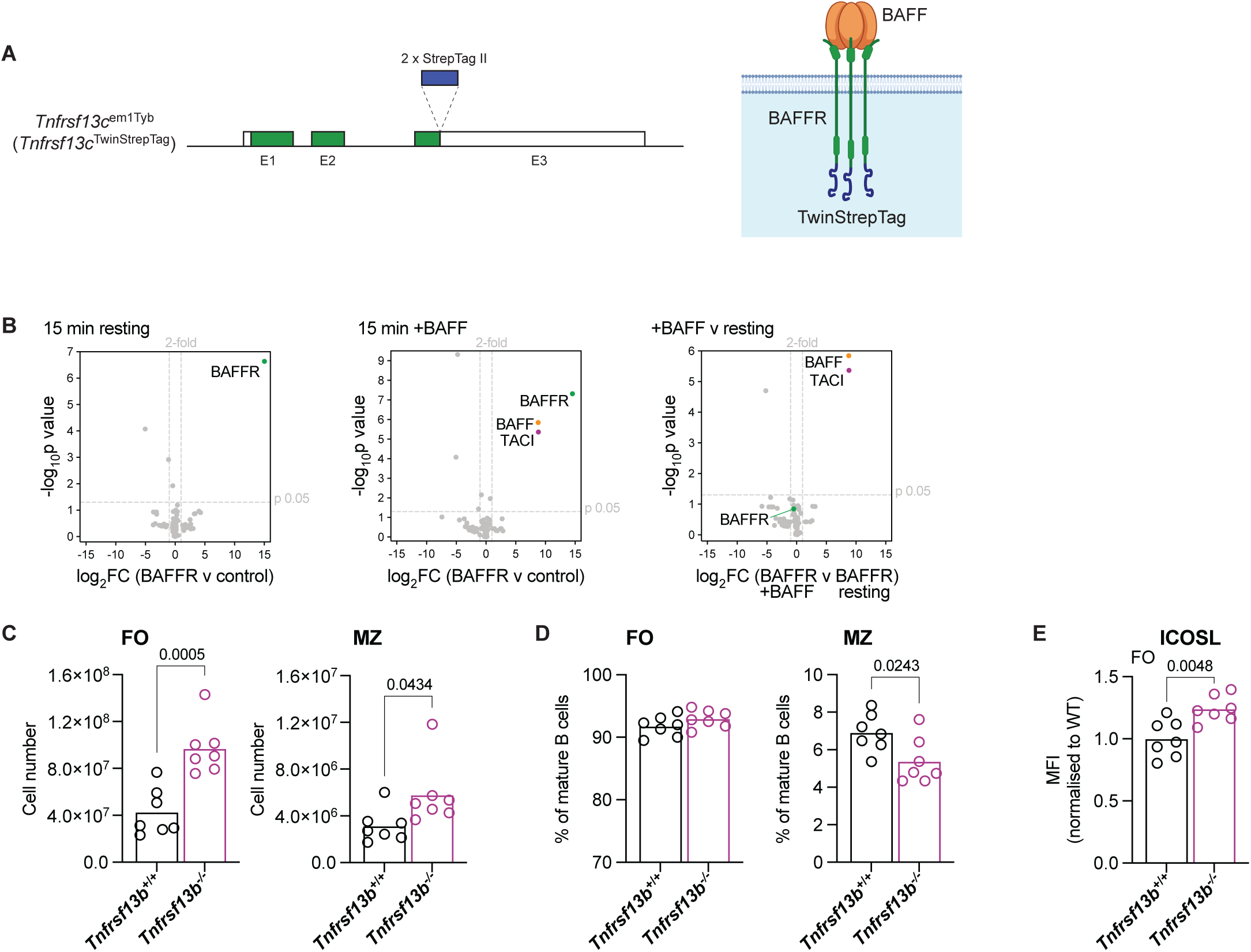
Analysis of the BAFFR interactome and TACI-deficient mice. A. Left, diagram of the *Tnfrsf13c*^em1Tyb^ allele showing exons 1-3 (E1-E3); filled boxes indicate coding sequence, open boxes indicate untranslated regions. Two StrepTag II sequences were inserted after the last coding codon in E3, before the stop codon. Right, diagram of a BAFFR- TwinStrepTag trimer showing the location of the affinity tag at the C-terminus of the protein. **B.** Volcano plots of proteins identified by mass spectrometry in BAFFR-TwinStrepTag affinity purifications (APs) from B cells activated with LPS, either resting or after stimulation with BAFF for 15 min. The plots show the log2-fold change (FC) in abundance of each protein in BAFFR APs compared to control APs from wildtype cells (log2FC(BAFFR v control)) or in stimulated BAFFR APs compared to resting BAFFR APs (log2FC(BAFFR +BAFF v BAFFR resting)) from three independent experiments, and the –log10p value determined by two-tailed Student’s t-tests. The thresholds used to determine specific BAFFR interactors (upper-right quadrant) are drawn at 2-fold enrichment and p = 0.05. The full list of proteins identified is in Table S1. **C, D.** The numbers (C) and proportions (D) of FO (CD1d^med^IgM^+^) and MZ (CD1d^hi^IgM^hi^) mature splenic B cells (CD93^-^B220^+^CD19^+^) in *Tnfrsf13b*^+/+^ and Tnfrsf13b^-/-^ mice. Bars show the mean of n = 7 mice, from 2 independent analyses. Each point represents one mouse. **E.** ICOSL surface expression, measured by flow cytometry, on splenic FO B cells (CD1d^med^IgM^+^CD93^-^B220^+^CD19^+^) from *Tnfrsf13b*^+/+^ and Tnfrsf13b^-/-^ mice, quantified as the geometric mean fluorescence intensity (MFI) and normalized to the mean MFI of *Tnfrsf13b*^+/+^ (WT) FO B cells within the same experiment. Bars show the mean of n = 7 mice, from 2 independent analyses. Each point represents one mouse. Statistical tests: unpaired t test. Numbers above graphs indicate p-values where p≤0.05 otherwise no value is shown.

**Figure S2.**
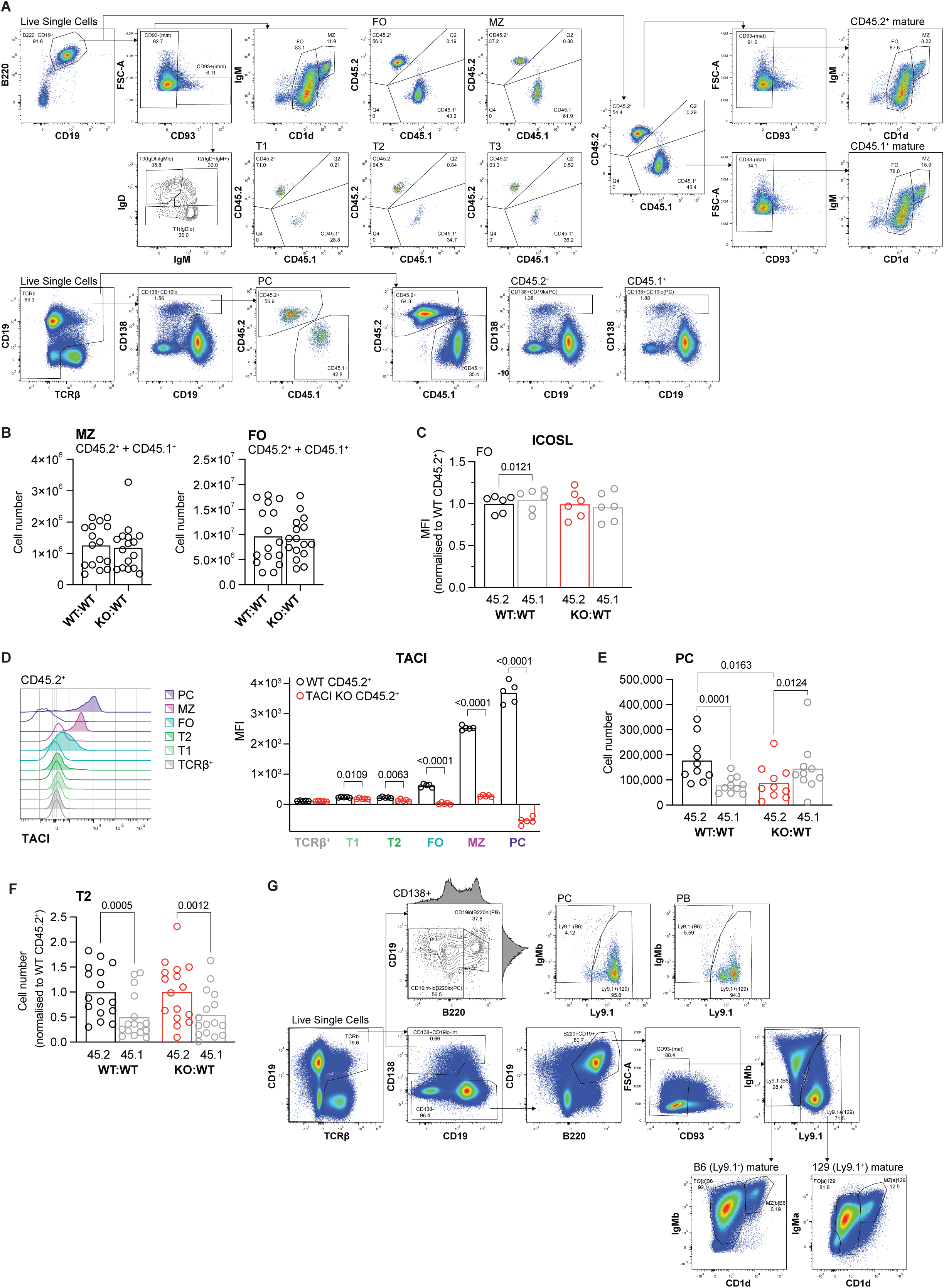
Analysis of mixed chimeras. **A.** Gating strategy used to identify FO, MZ, T1, T2, and T3 B cells and plasma cells (PCs) from the spleens of WT:WT and KO:WT chimeras. **B.** The total number of MZ (CD1d^hi^IgM^hi^) and FO (CD1d^med^IgM^+^) mature splenic B cells (CD93^-^ B220^+^CD19^+^CD138^-^; CD45.2^+^ and CD45.1^+^ combined) in WT:WT and KO:WT chimeras. Bars show the mean of n = 16 mice, from 4 independent analyses. Each point represents one mouse. **C.** ICOSL surface expression, measured by flow cytometry, on CD45.2^+^ and CD45.1^+^ splenic FO B cells (CD1d^med^IgM^+^CD93^-^B220^+^CD19^+^) from WT:WT and KO:WT chimeras, quantified as the geometric mean fluorescence intensity (MFI) and normalized to the mean MFI of WT CD45.2^+^ cells within the same experiment. Bars show the mean of n = 6 mice, from 2 independent analyses. Each point represents one mouse. Black: WT CD45.2^+^; red: TACI KO CD45.2^+^; grey: WT CD45.1^+^. **D.** TACI surface expression, measured by flow cytometry, on TACI KO and WT CD45.2^+^ splenic plasma cells (PC; TCRβ^-^CD138^+^CD19^lo^), MZ (CD93^-^CD1d^hi^IgM^hi^), FO (CD93^-^ CD1d^med^IgM^+^), T2 (CD93^+^IgD^+^IgM^+^) and T1 (CD93^+^IgD^lo^) B cells (B220^+^CD19^+^) and T cells (TCRβ^+^CD19^-^) from WT:WT and KO:WT chimeras. Left: representative flow cytometry histograms showing TACI expression on each cell type. Filled: WT; unfilled: TACI KO. Right: TACI expression quantified as the geometric mean fluorescence intensity (MFI). Bars show the mean of n = 5 mice, representative of 2 independent analyses. Each point represents one mouse. Black: WT CD45.2^+^; red: TACI KO CD45.2^+^. **E.** The numbers of CD45.2^+^ and CD45.1^+^ splenic plasma cells (PC; TCRβ^-^CD138^+^CD19^lo^) in WT:WT and KO:WT chimeras. Bars show the mean of n = 10 mice, from 2 independent analyses. Each point represents one mouse. Colors are as in C. **F.** The numbers of CD45.2^+^ and CD45.1^+^ splenic T2 (CD93^+^IgD^+^IgM^+^) B cells (B220^+^CD19^+^CD138^-^) in WT:WT and KO:WT chimeras. Bars show the mean of n = 16 mice, from 4 independent analyses. Each point represents one mouse. Colors are as in C. **G.** Gating strategy used to identify B6 (Ly9.1^-^) and 129 (Ly9.1^+^) FO B cells, MZ B cells, plasma cells (PC; CD138^+^B220^lo^CD19^int-lo^) and plasmablasts (PB; CD138^+^B220^hi^CD19^int^) from the spleens of WT:wt and KO:wt chimeras, for the data shown in Figure 2. Statistical tests: unpaired t test (B), repeated measures two-way ANOVA with Fisher’s LSD test (C, E, F), unpaired Welch’s t test corrected for multiple comparisons using the Holm-Šidák method (D). Numbers above graphs indicate p-values where p≤0.05 otherwise no value is shown.

**Figure S3.**
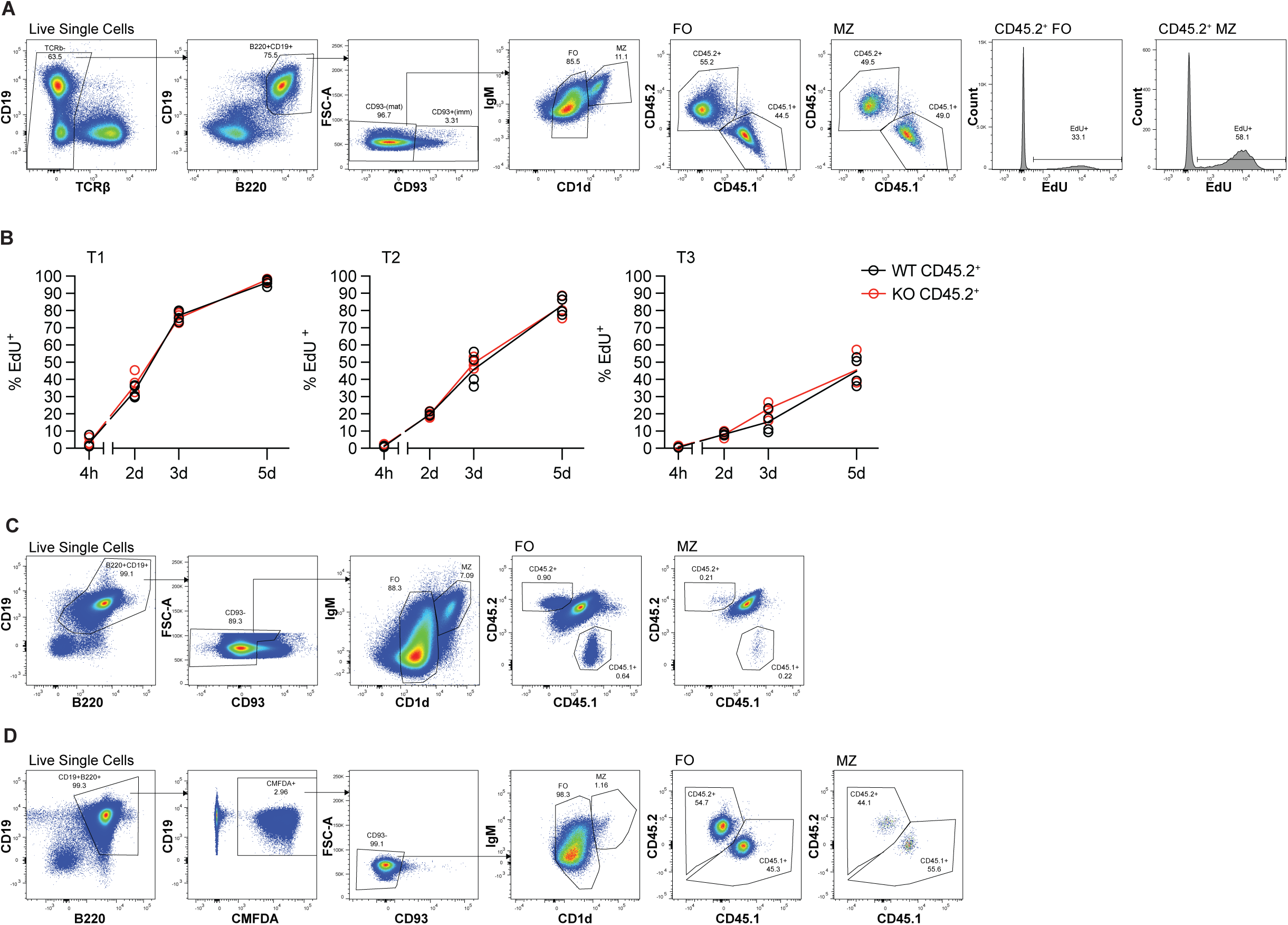
Analysis of EdU labeling and adoptive transfer studies. **A.** Gating strategy used to identify EdU^+^ CD45.2^+^ FO and MZ B cells in splenocytes from WT:WT and KO:WT chimeras after continuous EdU labelling *in vivo*. **B.** The proportion of EdU^+^ cells within T1 (IgD^lo^), T2 (IgD^+^IgM^+^) and T3 (IgD^hi^IgM^lo^) CD45.2^+^ immature splenic B cell populations (CD93^+^B220^+^CD19^+^) from WT:WT and KO:WT chimeras at 4 h, 2 days, 3 days and 5 days after EdU injection (performed at 12 weeks post-reconstitution) and EdU continuously administered via drinking water. Each point represents one mouse (4 h WT n = 4, KO n = 5; 2 d, 3 d and 5 d n = 4). Lines connect the mean at each timepoint. Black: WT CD45.2^+^; red: TACI KO CD45.2^+^. No difference reported between WT and KO means at any timepoint as determined by two-way ANOVA with Šídák’s multiple comparisons test. **C.** Gating strategy used to identify transferred CD45.2^+^ and CD45.1^+^ FO and MZ B cells in the spleens of CD45.2^+^CD45.1^+^ recipients, 7 days post-adoptive transfer. **D.** Gating strategy used to identify transferred CMFDA-labelled CD45.2^+^ and CD45.1^+^ FO and MZ B cells in the spleens of CD45.2^+^CD45.1^+^ recipients, 1 h after adoptive transfer.

**Figure S4.**
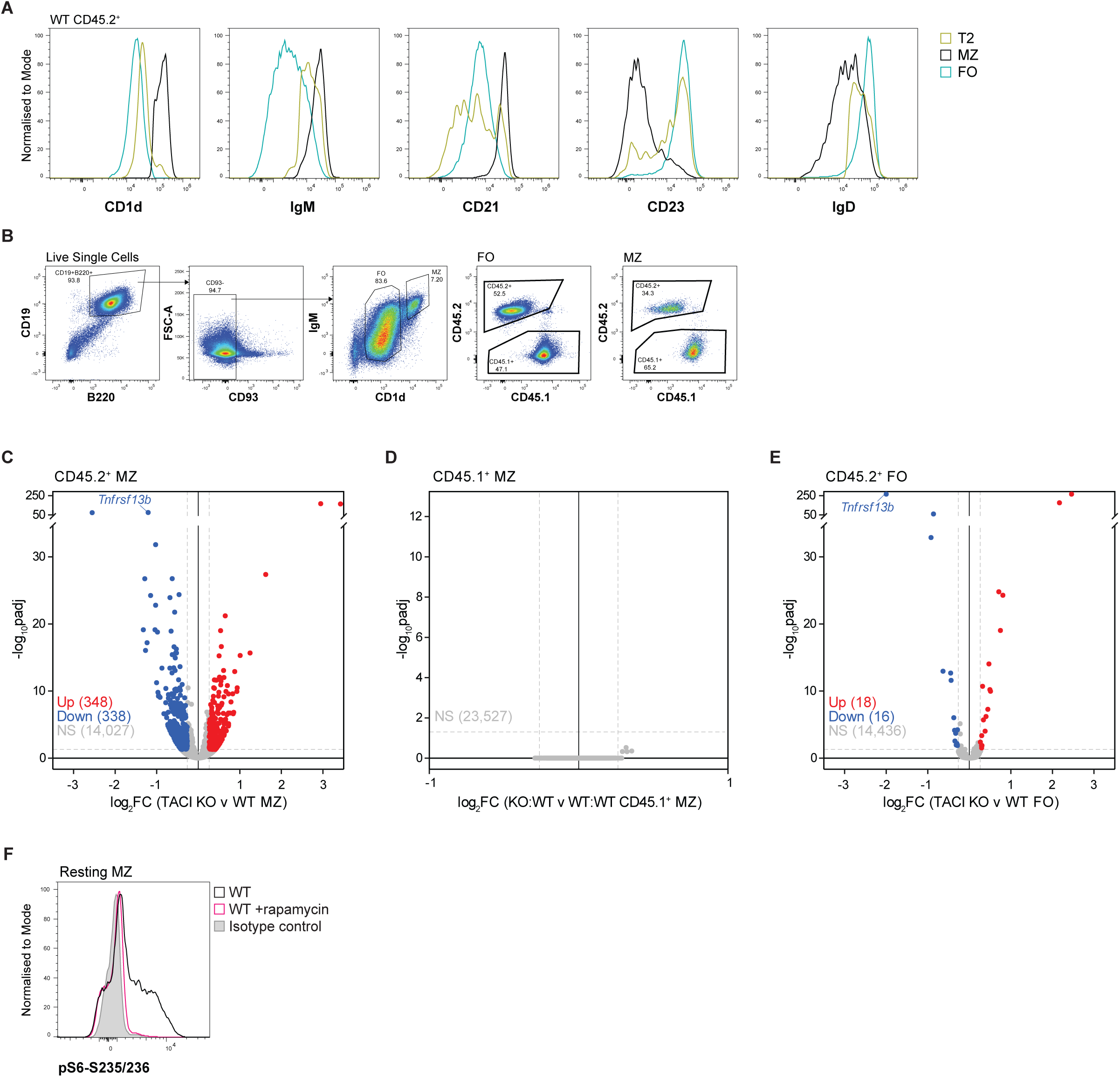
Surface marker expression, RNAseq analysis and pS6 staining. **A.** Flow cytometry histograms comparing CD1d, IgM, CD21, CD23 and IgD surface expression on T2 (IgD^+^IgM^+^CD93^+^), MZ (CD1d^hi^IgM^hi^CD93^-^) and FO (CD1d^med^IgM^+^CD93^-^) WT CD45.2^+^ B cells (B220^+^CD19^+^) from the spleens of WT:WT chimeras. Gold: T2; black: MZ; blue: FO. **B.** Gating strategy used to sort CD45.2^+^ and CD45.1^+^ FO and MZ B cells from the spleens of WT:WT and KO:WT chimeras by FACS for RNA sequencing. **C-E.** Volcano plot of differential gene expression between TACI KO and WT CD45.2^+^ MZ B cells (C), between WT control CD45.1^+^ MZ B cells (D) and between TACI KO and WT CD45.2^+^ FO B cells (E) from KO:WT and WT:WT chimeras, respectively, showing the mean log2- foldchange (log2FC) and the -log10 adjusted p-value (-log10padj) for each gene from n = 6 mice. Significantly differentially expressed genes are in red (upregulated; log2FC ≥ 0.263, padj < 0.05) and blue (downregulated; log2FC ≤ -0.263, padj < 0.05). Grey: not significantly differentially expressed (NS). Dashed grey lines show the thresholds for 1.2-fold change in gene expression and padj = 0.05. **F.** Flow cytometry histograms comparing pS6-S235/236 levels in resting WT MZ B cells at 37°C with those treated with the mTORC1 inhibitor rapamycin. Black: WT MZ; pink: WT MZ +10 nM rapamycin; grey: WT MZ cells stained with isotype control antibody.

